# Identification and characterization of thousands of bacteriophage satellites across bacteria

**DOI:** 10.1101/2022.09.14.508007

**Authors:** Jorge A. Moura de Sousa, Alfred Fillol-Salom, José R. Penadés, Eduardo P.C. Rocha

## Abstract

Bacteriophage-bacteria interactions are affected by phage satellites, elements that exploit phages for transfer between bacterial cells. Satellites can encode defense systems, antibiotic resistance genes, and virulence factors, but their number and diversity are unknown for lack of a tool to identify them. We developed a flexible and updateable program to identify satellites in bacterial genomes – SatelliteFinder – and use it to identify the best described families: P4-like, phage inducible chromosomal islands (PICI), capsid-forming PICI, and phage-inducible chromosomal island-like elements (PLE). We vastly expanded the number of described elements to ∼5000, finding hundreds of bacterial genomes with two different families of satellites, and dozens of *Escherichia coli* genomes with three. Most satellites were found in Proteobacteria and Firmicutes, but some are in novel taxa such as Actinobacteria. We characterized the gene repertoires of satellites, which are variable in size and composition, and their genomic organization, which is very conserved. With the partial exception of PICI and cfPICI, there are few homologous core genes between families of satellites, and even fewer homologous to phages. Hence, phage satellites are ancient, diverse, and probably evolved multiple times independently. Occasionally, core genes of a given family of satellites are found in another, suggesting gene flow between different satellites. Given the many elements found in spite of our conservative approach, the many bacteria infected by phages that still lack known satellites, and the recent proposals for novel families, we speculate that we are at the beginning of the discovery of massive numbers and types of satellites. SatelliteFinder is accessible for the community as a Galaxy service at https://galaxy.pasteur.fr/root?tool_id=toolshed.pasteur.fr/repos/fmareuil/satellitefinder/SatelliteFinder/0.9

## Introduction

Bacteriophages (phages) shape the evolution and composition of bacterial communities, both through predation and by driving horizontal gene transfer between bacterial cells (1, 2). Phages are notorious parasites of bacteria and are themselves parasitized by phage satellites. These elements lack some of the functions required for autonomous horizontal transfer, which they hijack from helper phages. In the past, phage satellites were sometimes mistaken for defective phages, even if their gene repertoires rarely have genes in common with phages. Yet, some phage satellites do have phage-like genes. For example, the phage-inducible chromosomal islands (PICI) of *Staphylococcus aureus* typically encode packaging functions homologous to those of the helper phage 80α (3, 4). In contrast, the P4 satellite of *Escherichia coli* has few homologs with its P2 helper phage (5). Phage satellites exploit their helper phages through molecular mechanisms that depend on the type of satellite. The extent of this exploitation is variable, as are its consequences for phage reproduction. Phage-inducible chromosomal island-like elements (PLE) of *Vibrio cholerae* completely block the propagation of the helper lytic phage ICP1 (6) and PICI can severely reduce phage reproduction (7). On the other hand, the recently identified capsid-forming PICI (cfPICI) EcCIEDL933 of *E. coli* has negligible effects on phage fitness (8). Beyond their effect on phage infection, satellites provide their bacterial hosts with accessory potentially adaptive functions. For example, some PICI encode virulence factors, like toxins, or antibiotic resistance genes (3), and some P4-like elements and PICI encode anti-phage immunity systems (9, 10). Hence, satellites have wide functional and ecological impacts in phages and in bacteria.

The few satellites that have been studied in detail encode a set of core functions that are sometimes non-homologous but can be grouped into four major groups: integration, regulation, replication and hijacking. All known satellites are integrated in chromosomes, which usually occurs by the action of an integrase of the Tyrosine recombinase family. Upon excision, satellites require specific replicases to replicate before being packaged in viral particles. Genetic regulation is essential for the success of the element. On one hand, the element may remain for long periods of time largely silent in the chromosome before entering in contact with an infecting helper phage. On the other hand, upon co-infection by a helper phage, the satellite must coordinate its expression with the latter. One of the most fascinating traits of satellites is the diversity of mechanisms they use to subvert the host phage viral particles. Some phage satellites physically constrain the size of the viral capsid produced by the phage, so that phage DNA does not fit inside – but the satellite DNA does (11). Other satellites directly redirect the packaging of their own DNA into the viral capsid by encoding sequences that mimic the phages’ terminases (12).

Phage satellites have only recently emerged as a distinct class of mobile elements. Even if the satellite P4 was discovered decades ago, few phage satellites have been detected in genomes until very recently. Also, some of these mobile elements were either mistaken as plasmids (13) or annotated as defective phages or prophage-like remains (14). However, this perspective has recently changed. There is increasing evidence of the pervasiveness and importance of phage satellites (15, 16), and recent work has started to uncover the diversity of these elements, especially within a given family (5, 12). These studies have helped recognize phage satellites as characteristic mobile elements that have specialized in being mobilized by fully functional phages. Moreover, there have been recent reports of genomic islands transduced by phages that are different from known phage satellites, which suggests that different types of satellites remain to be discovered (17, 18).

Many studies have unraveled the mechanisms of function of the model satellites for each of the known families, as well as their importance in bacterial evolution and pathogenicity (4, 9, 10, 19). But their abundance in genomes is poorly studied for lack of a systematic way to identify them. This is important, because the small number of known phage satellites has limited their comprehensive study in terms of evolution and diversity. Recently, we have reported the discovery of ca 1000 elements of the P4-like family, which has led to novel insights regarding their diversity, evolution, genomic composition and organization (5). Here, we systematize and expand this analysis to all currently known phage satellite families and report the discovery of more than 5000 putative phage satellites in complete bacterial genomes. This allowed us to study the abundance of phage satellites within and across bacterial hosts, and to understand how the different families of satellites are organized in terms of their core components and genetic repertoires. We also sought to understand whether there are similarities between the different main families of phage satellites, and whether there are different sub-families within them. Our approach allows for a novel, automatic detection of phage satellites, of all known families, in bacterial genomes and sheds light on the abundance and diversity of these mobile elements.

## Methods

### Genomic datasets

We retrieved all the complete genomes of the NCBI non-redundant RefSeq database (ftp://ftp.ncbi.nlm.nih.gov/genomes/refseq/, last accessed in March 2021), including 21086 bacteria, 21520 plasmids and 3725 phages. We also retrieved and syntactically annotated both complete and draft genomes of *Vibrio* spp (n=11627 in total), using PanACoTA (version 1.3.1 (20)), ran with the Singularity module. We used the methods “prepare -s <species>“ (where <species> was iteratively replaced with the most representative species of Vibrionacea: Vibrio, Aliivibrio, Enterovibrio, Photobacterium, Grimontia, Salinivibrio and Thaumasiovibrio, databases accessed in March 2022), with the options “--min 0 --max 0.4”, for the MASH distance thresholds (21). We then used the method “predict --prodigal” to syntactically annotate the genomes.

### Overall strategy for detection of phage satellites

There are only a few experimentally verified satellites. This means that we cannot use machine learning methods to identify the elements at this stage. Instead, we made a curated annotation of a large set of known satellite elements and used them to iteratively identify core genes of each family of satellites. We then designed individual and customizable MacSyFinder (22, 23) models that represent the genetic composition of the different phage satellite families (**Fig S1**). The models were used in MacSyFinder to identify occurrences of the putative satellites. MacSyFinder missed a few elements in certain genomic contexts or was unable to disentangle tandem occurrences of similar satellites. Hence, to improve the methods, we developed a post-processing script, which also provides a classification of the satellite variants (**Fig S2**).

Throughout the process of optimizing the detection of satellites, we sometimes identified components that are less conserved than expected and were thus removed from the list of core genes. Also, some genes were found to be more conserved than expected and were included as core components. As a result, we iterate the entire process again, until very few changes occur between iterations. At each iteration we verify that most of the known satellites are accurately classified. In the following sections, we describe the key steps of the process that leads to our method of detecting phage satellites. The details associated with each family of satellites are given in the initial paragraphs of each section of the results, and the tables with all the satellites identified are given in **File S1**.

### Modelling of satellite-like systems

We made an iterative enrichment and curation of the core (marker) components based on the analysis of genomic regions in bacterial genomes that could correspond to putative satellites.

1. We collected genomic regions containing the pre-defined components at less than a pre-defined distance, to minimize the detection of components of tandem elements. For example, two components are clustered if they are less than 10 genes apart. We make a transitive clustering of the components, i.e., when one advances linearly in the genome sequence the cluster is closed and evaluated only when meeting a succession of more than 10 genes lacking a marker. If the cluster contains enough marker genes, i.e. it meets the minimal quorum, it is kept for further analysis. We checked at this stage that the procedure identified the known or previously proposed satellites and does not match known phages.
2. We took these clusters and built their pangenome to identify the most abundant gene families in each phage satellite family. The pangenomes were built by clustering all the candidate proteins at a minimum of 40% identity, using mmseqs2 (24) (version 13-45111), with parameters –cluster_mode 1 and –min_seq_id 0.4 (all other parameters were left as default). The clusters of abundant gene families are usually good markers for the presence of a satellite.
3. The resulting gene families were functionally annotated using PFAM (release 33.1) and bactNOG (25).
4. Sometimes a family that was not initially used as a marker was found to be present at high frequency in #2. In such cases, we checked if an adequate HMM profile was available in PFAM. If this was not the case the HMM profile was built as described above. In any case, the inclusion of novel profiles implicated re-starting the process (back to #1).
5. We tested if we should define groups of marker genes as “exchangeable”. In such cases, MacSyFinder will fill the quorum of a marker if it identifies one of a set of protein profiles. For example, integrases of the Tyrosine recombinase and of the Serine recombinase families are often found as functional analogs in mobile genetic elements. To identify these exchangeable elements, we used the literature (for analogs) or searched the known satellites for analogous components. Introduction of “exchangeable” components led us back to #1.
6. We varied the parameter of maximal distance between consecutive components to test how it affects the method. When the analysis revealed that we should extend this distance because some frequent gene families were often found a bit further downstream, we adapted this parameter. If this increased the frequency of a gene family up to the point where it could be used as a marker, we started again at #1. In the cases presented in this study, a distance of 10 or 15 genes was regarded as a good trade-off between identifying a few unusually long elements and preventing the frequent aggregation of several satellites into one cluster.
7. Sometimes we added components in the models to improve the discrimination between different families of satellites, or between satellites and prophages. In most cases, this was done by including some “forbidden” components that are almost never found in one family of satellites. For example, tail proteins are never (or very rarely) found in satellites and allow to distinguish them from prophages. When such markers were added, we started again at #1.

This iterative procedure resulted in a MacSyFinder model for each family of satellites. These models were used to systematically search for putative phage satellites in bacterial genomes. The MacSyFinder models for each satellite family are available within the Docker package of SatelliteFinder (https://hub.docker.com/r/gempasteur/satellite_finder).

### Identification of markers and construction of HMM profiles

The first step in identifying novel satellites is to characterize the known ones and identify the genes that are most frequently associated with them (markers). We collected the satellites experimentally verified or proposed as homologs in previous publications and clustered them by 40% (or 20%, for cases where no clusters emerged) protein sequence similarity. This revealed families of proteins that were the most frequent in a given family of satellite. The majority of these frequent components was adequately identified by existing HMM profiles of the PFAM database. When this was not the case, we built custom HMM profiles by aligning all sequences of the family with Clustal Omega (26) (Version 1.2.3) with default parameters, and then by using hmmbuild (default parameters) from hmmer 3.3.2 (27).

### Identification of putative phage-satellites with MacSyFinder

We used MacSyFinder (22, 23) (version 2.05rc) to provide a reproducible, shareable, and easily modifiable tool to identify phage satellites in bacterial genomes. Briefly, MacSyFinder searches for co-occurrence of the markers of each phage satellite family in bacterial genomes. The criteria for the identification of the occurrences of markers and for the acceptable patterns of co-occurrence can be defined by the user and that is what we call a model. MacSyFinder reports the cases with highest scores, namely where the largest co-occurrence of the different markers was found. For example, a genomic region with all markers gets a higher score than a genomic region with fewer markers.

Typically, one searches for these markers either using curated thresholds or by providing an e-value cutoff. Here, we lacked previous information on protein sequence diversity in the satellites and we tried to maximize the sensitivity of the models relative to this parameter. Hence, we used general and relaxed cutoffs (e-value < 0.01 and coverage > 40%, with parameters “--no-cut-ga --i-evalue-sel 0.01 --coverage-profile 0.4” in MacSyFinder), to detect distant homologs. The MacSyFinder model requires the identification of a number of markers and their co-occurrence. The need to respect a minimum quorum of co-localized components decreases the rate of false positives that could arise from the use of relaxed criteria of sequence similarity, because false positive satellites would require the random simultaneous co-localization of individual false positive markers. A specific concern arises with degraded prophages that could resemble some satellites. These are discussed in the corresponding results section. The remaining parameters of MacSyFinder were left as default.

### Post-processing of MacSyFinder results

The use of MacSyFinder alone allows to find most of the known or previously proposed satellites (**Table S1**). Yet, we noticed that a few elements were lost in some families (e.g., 10% of cfPICI elements). This is caused by two features of the program that we corrected by post-processing the results.

1. Clusters of genes with a match to at least one forbidden profile are rejected. This feature is required to distinguish satellites with markers often found in phages from phages themselves, or to distinguish between different satellite families with many homologous core components. However, sometimes satellites and prophages are contiguous in bacterial genomes and the forbidden component is on the flanking element, not on the satellite itself. Hence, we post-process the results to include those discarded due to the presence of one single “forbidden” component in the cluster (to allow for unknown variants) and those where the “forbidden” component is outside of the cluster of components (prophages contiguous to satellites). These “rescued” clusters are very rare for most satellite types (see Results section). They are specifically identified in the output of our scripts and in our analyses in the main text.
2. MacSyFinder outputs the largest possible cluster of markers of satellites, which may result in merging multiple satellites. It may also merge satellites with small contiguous mobile elements having an integrase. Our post-processing starts by handling the occurrence of multiple integrases and then focuses on tandem satellites. First, we search for the presence of multiple integrases (we assume there should be only one). If so, we choose the one that is closest to the other (non-integrase) components, and use any other integrase as a starting point of a new set of components. This procedure may thus output several putative satellites from a single MacSyFinder cluster.

The MacSyFinder output lists the markers present in each putative satellite. For the analysis of the number and types of components present in the latter, we post-process the output to classify putative satellites into “types”: (A) have all (N) core components, (B) have N-1 core component, (C) have N-2 core components, and so on. These types are further categorized as “variants” that correspond to the component(s) that are missing. We note that many Type B and C elements are complete and functional, since some correspond to elements that were experimentally verified and many are very conserved. They may correspond to a variant of the prototypical satellite that either completely lacks a given marker component (e.g., due to gene loss or pseudogenization), has a non-homologous analog of the component, or has a diverged component that was not detected.

### Genomic comparison of phage satellites with weighted gene-repertoire relatedness (wGRR)

We searched for sequence similarity between all proteins of phages and/or satellites using mmseqs2 (24) with the sensitivity parameter set at 7.5. The results were converted to the blast format and we kept for analysis the hits respecting the following thresholds: e-value lower than 0.0001, at least 35% identity, and a coverage of at least 50% of the proteins. The hits were used to retrieve the bi-directional best hits between pair of genomes, which were used to compute a score of gene repertoire relatedness weighted by sequence identity (28):

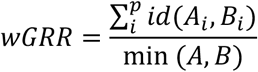

where A_i_ and B_i_ is the pair *I* of homologous proteins present in *A* and *B*, id(*A*_*i*_,*B*_*i*_) is the sequence identity of their alignment, and min(*A,B*) is the number of proteins of the genome encoding the fewest proteins (*A* or *B*). wGRR is the fraction of bi-directional best hits between two genomes weighted by the sequence identity of the homologs. It varies between zero (no bi-directional best hits) and one (all genes of the smallest genome have an identical homolog in the largest genome). wGRR integrates information on the frequency of homologs and sequence identity. For example, when the smallest genome has 10 proteins, a wGRR of 0.2 can result from two homologs that are strictly identical or five that have 40% identity. The hierarchical clustering of the wGRR matrix, and the corresponding heatmap, were computed with the *clustermap* function from the *seaborn* package (version 0.11.1, developed for Python 3.9), using the Ward clustering algorithm.

### Calculation of diversity of bacterial hosts

We assessed the diversity of the phage satellites’ bacterial hosts using two measures. The Species Diversity (S) is the number of bacterial species where at least one phage satellite was identified. Since the genome datasets are very unbalanced, with some species accounting for a large fraction of genomes, the Species Diversity measure can misrepresent the true diversity of the hosts of satellites. Hence, we use also Shannon’s diversity index (H’) (29) for each phage satellite family. The index is calculated according to the formula:

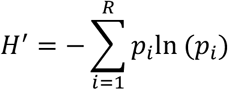

Where R is the total number of bacterial species with at least one satellite, and *p*_*i*_ is the proportion of satellites found in the ith bacterial host species, in relation to the total number of satellites for the corresponding family (e.g., if there is a total of 500 elements, and 300 are found in *E. coli* genomes, *p*_*E*.*coli*_=0.6). Thus, a family of satellites where the vast majority of elements is located in only a few bacterial species will have a lower Shannon diversity index than a family of satellites that is equally spread across bacterial species.

### Phylogenetic analysis

We aggregated in single fasta files all the protein sequences corresponding to either the regulatory, capsid or small terminase components of both PICI and cfPICI genomes. The sequences were aligned using mafft-linsi (30) (v. 7.490, default parameters) and the resulting alignment trimmed with clipkit (31) (v. 1.3.0, default parameters). We then used IQ-Tree (32) (v. 1.6.12) to build the phylogenetic trees, with the options –bb 1000 to run the ultrafast bootstrap option with 1000 replicates and –nt 6. The resulting tree files were visualized and edited using the v5 webserver of iTOL (33).

#### Availability

We provide the MacSyFinder models and the customized python scripts to perform the abovementioned post-processing as a tool we call SatelliteFinder, that is available as a Docker package (https://hub.docker.com/r/gempasteur/satellite_finder) and also a Galaxy server interface (34) (https://galaxy.pasteur.fr/root?tool_id=toolshed.pasteur.fr/repos/fmareuil/satellitefinder/SatelliteFinder/0.9).

## Results

### P4-like satellites are frequently found in Enterobacterial genomes

The P4 satellite is among the best studied satellites (11, 35, 36). P4 hijacks P2 capsids through physical constraining, in order to encapsidate its own DNA. Our previous work has shown that the P4-like family of satellites contains seven very conserved components (5). We used this information to build MacSyFinder models to detect P4-like satellites (see Methods). We searched for the co-occurrence of an integrase with six other components: Psu, Delta and Sid, which are involved in the hijacking of the capsid of the P2 helper phage; a regulatory protein, typically homologous to AlpA (although we also search for homologs of MerR or Stl as other possible regulators); Ash (also called ε), which inactivates the repressor of the helper phage, causing its induction; and α, a protein with primase and helicase activities that is required for P4 replication.

The P4 MacSyFinder model identified a similar amount (1054 vs 1037) of P4-like elements relative to our previous search in the same database (5). Around 90% of the elements were in the same bacterial host genome and started at the same integrase gene. We used this model to search for P4-like elements in a much larger dataset of 21086 bacterial genomes (**Fig 1**). We more than doubled the number of putative P4-like satellites previously identified (2160). The majority (1621) of these elements encode all the seven core components (henceforth called Type A, **Fig 1A and 1B**), whilst 350 elements lacked one of them (Type B). The missing component may be absent, be non-identifiable by the protein profile, or may be replaced by a functional analog (see Methods). The most abundant of these variants lacks Psu (Type B#var06). Since Sid and Psu are structural homologs (37) it is possible that some variants of the former may compensate for the absence of the latter. There are 189 elements that lack two core components (Type C), most often ε and α (Type C#var01) or AlpA and α (Type C#var02). The vast majority of putative P4-like phage satellites (93%) were detected in Enterobacteriaceae, where 35% of the bacterial genomes encode from one to three of these elements (**Fig 1C and 1D**). Other bacterial families with P4-like elements include Yersiniaceae (23% of the genomes with at least one element), where variants lacking Psu are prevalent, Pectobacteriaceae, Erwinaceae and Hafniaceae. All but one P4-like elements are integrated in bacterial chromosomes, confirming that these elements are usually not present in cells as plasmids.

**Figure 1.**
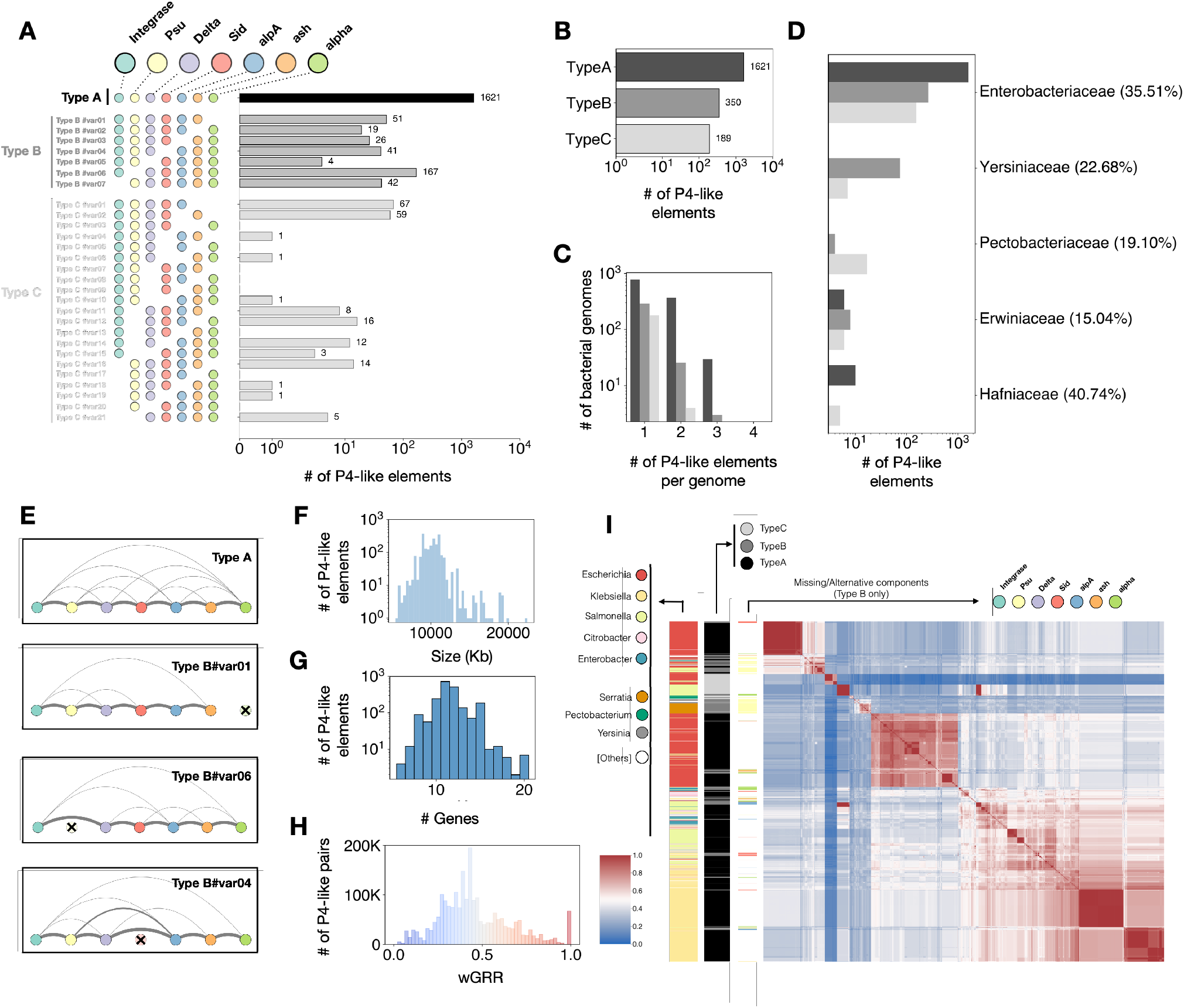
The abundance, genetic organization, bacterial hosts and genomic structure of the family of P4-like elements. **A)** Number of the different variants of P4-like elements identified in bacterial genomes. **B)** Total number of the different types of P4-like elements. **C)** Total number of P4-like genomes per bacterial genome. **D)** Distribution of P4-like elements in bacterial families. The percentages in front of each family correspond to the proportion of genomes of that family where (at least) one P4-like element was inferred. **E)** Genomic organization of the four most frequent variants of P4-like satellites. Nodes correspond to the different markers of the satellite (variants where the marker is absent have that indicated as a crossed-out node) and the edges between the nodes represent the frequency with which those two components are contiguous (but not necessarily adjactent). **F)** Distribution of the sizes of the extracted genomes of P4-like elements. **G)** Distribution of the number of proteins contained within each P4-like genome, for the elements detected in the two different bacteria datasets. **H)** Distribution of the pairwise wGRR values between all the P4-like genomes. **I)** Symmetric heatmap of the matrix of the wGRR values ordered using hierarchical clustering. The colours follow the same code as in (**H**) with blue pixels representing low wGRR values (dissimilar genomes) and red pixels representing high wGRR values (similar genomes). The columns to the left of the heatmap indicate the bacterial species where the P4-like genome was detected, the Type (A, B or C) of the P4-like genome, and the component that is missing for given variants (exclusively for elements of Type B).

Consistent with our previous analysis, the organization of the core components of P4-like satellites is conserved. Type A and the most frequent Type B variants encode the *psu*-*delta*-*sid* operon, followed by *alpA*, ε and α (**Fig 1E** and **Fig S3A**), with each component usually found in the same relative location (**Fig S3B)**. We delimited the element between the integrase and its farthest core component, discarding those lacking an integrase. The resulting 2097 P4-like elements have a median size of 10Kb and a median of 11 genes (**Fig 1F, Fig 1G**). A small minority of these (<2%) is larger (between 15 and 22Kb).

We measured the similarity between P4-like elements using weighted gene repertoire relatedness (wGRR, see Methods). Some elements are identical (peak at wGRR=1 in **Fig 1H**), but most of them are only moderately related. The median wGRR is 0.42. We clustered them in relation to wGRR to assess their similarities (**Fig 1F**). There are two distinct sub-families within *Escherichia* genomes, and an additional large family including mainly elements found in *Salmonella* and *Klebsiella* genomes, but also in *Enterobacter* and *Citrobacter*. Other subfamilies include other clades such as *Serratia, Yersinia* and *Salmonella*. The subgroup of elements specific to *Salmonella* are all of Type C, as are the small family of elements in *Escherichia* genomes that form a very distinctive subfamily at the top of the matrix in **Fig 1F**. Many elements lack the same core genes and are associated with specific clades. This suggests that these are not defective elements. Instead, they seem to form distinctive variants (or subfamilies) of the P4 family. If the functions of the missing core genes in these variants are facultative, or if they can be complemented by other components remains unknown.

Although subfamilies tend to be associated with specific bacterial hosts, we also found some very similar elements in different species. This suggests of horizontal transfer of P4-like satellites across distant bacteria. We found 3182 pairs of very similar elements in different bacterial species (4% of those with wGRR≥0.9). Some very similar (wGRR > 0.8) pairs can also be found in different families (280 pairs, e.g., between Enterobacteriaceae and Hafniacea, or Erwiniaceae). Together, these results reinforce our previous findings of a large (and now even larger) family of P4-like phage satellites with a characteristic and conserved genomic organization and a broad host range.

### Phage Inducible Chromosomal Islands (PICI) are diverse and widespread across bacterial phyla

PICI include the *Staphylococcus aureus* pathogenicity islands (SaPI) that were extensively studied, as well as numerous other elements present in both diderm and monoderm bacteria (12, 38). The described PICI have a conserved genetic organization and five core components found in almost all known elements: (i) integrase, (ii) regulation module (homolog to *alpA, merR* or *stl*), (iii) primase-replicase module, (iv) capsid morphogenesis (more frequent in PICI from Proteobacteria), encoding a protein that is thought to modify the morphology of the hijacked capsids to block the encapsidation of phage DNA, (v) small terminase subunit, responsible for redirecting the packaging of the capsid to the satellite’s DNA. While other accessory genes are normally encoded by PICI (notably between the integrase and the regulation or the primase-replication module, or after the *terS* homolog), PICI typically do not encode other phage-like structural or lysis genes. We used the core components to detect PICI (**Fig S1B**). Given the homologs between PICI, cfPICI and prophages, we included in the PICI model several “forbidden” components to discriminate them accurately from the others (see Methods): tail, holin, and cfPICI components described below.

Our approach identified 1436 putative PICI, the vast majority (>99.9%) in chromosomes (**Fig 2A and B**): 375 (26%) with the five core components (Type A), and 1061 (74%) with four (Type B). We note that a small number of them (31, 2%) were initially rejected by MacSyFinder due to the presence of one forbidden gene near (but outside) the cluster of PICI-like core components. They were recovered by post-processing the results (see Methods). The vast majority (97%) of the elements with all core components (Type A) are found in *Escherichia coli* genomes. The other variants are found across much more diverse bacterial hosts (**Fig 2D**), including in 66% of Mycobacterial genomes (including *M. tuberculosis*) and 35% of Staphylococcaceae genomes. Some of the latter were previously identified as SaPIs. They are known to lack the capsid morphogenesis gene typically found in Enterobacterial PICI because they rely in a different hijacking strategy (3). Thus, these variants are *bona fide* functional PICI, experimentally observed to be mobilized by helper phages. We also detected more than 6000 elements with three core genes (Type C). These may be functional satellites, but the small number of core genes increases the probability that they may be defective PICI, other mobile genetic elements, or just random aggregates of PICI-like functions (for instance, a typical P4-like satellite is classified as a Type C PICI because it encodes an integrase, AlpA and a primase/replicase). Hence, in order to remain conservative in our analysis, we focused on PICI-like elements of Type A and B.

**Fig. 3.**
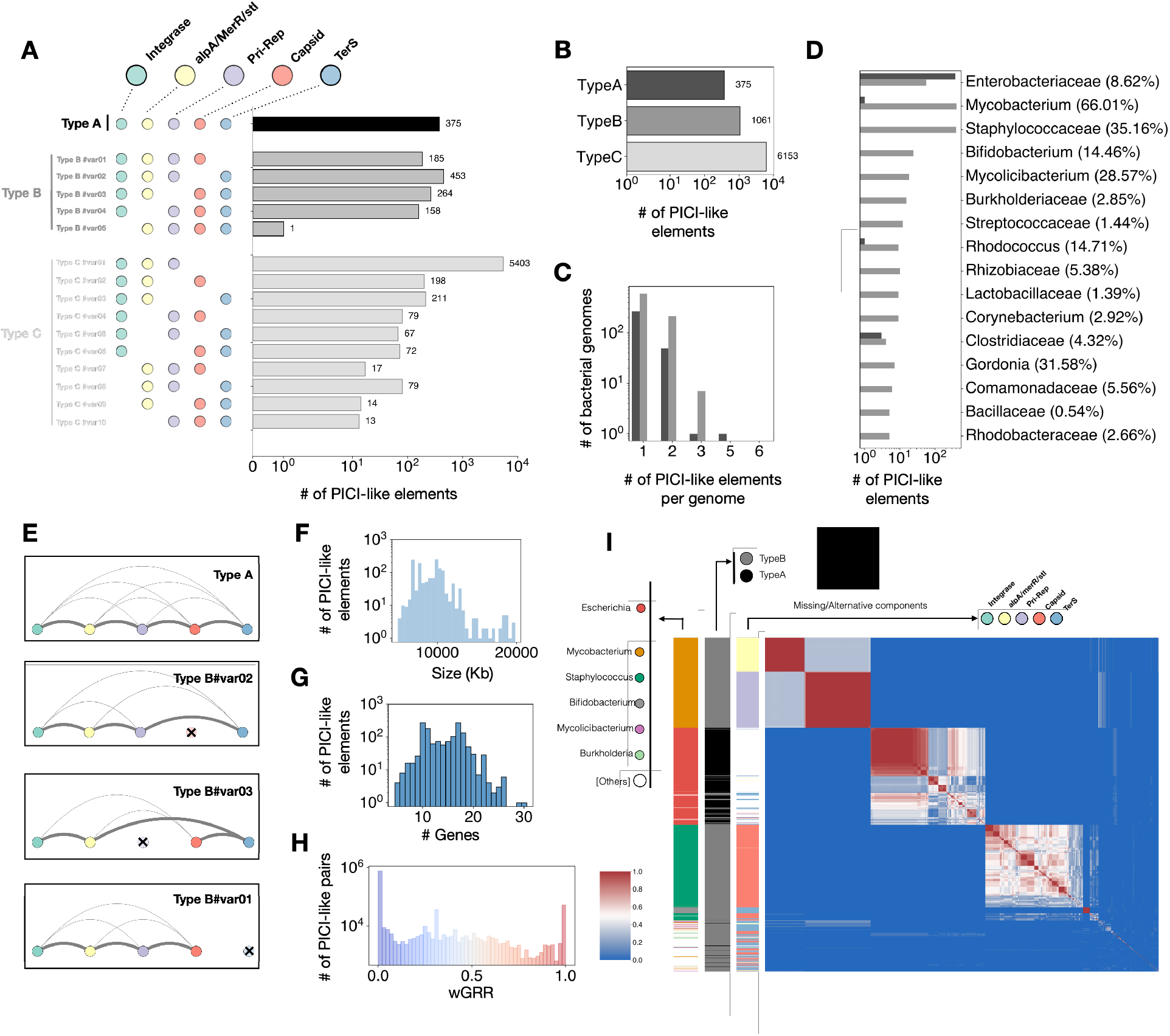
The abundance, genetic organization, bacterial hosts and genomic structure of the PICI family. **A)** Number of the different variants of PICI-like elements identified in bacterial genomes. **B)** Total number of the different types of PICIs. **C)** Total number of PICIs per bacterial genome. **D)** Distribution of PICIs of Type A or B in bacterial families. The percentages in front of each family correspond to the proportion of genomes of that family where (at least) one PICI was inferred. **E)** Genomic organization of the four most frequent variants of PICIs. Nodes correspond to the different markers of the satellite (variants where the marker is absent have that indicated as a crossed-out node) and the edges between the nodes represent the frequency with which those two components are contiguous (but not necessarily adjacent). **F)** Distribution of the sizes of the extracted genomes of PICIs. **G)** Distribution of the number of proteins contained within each PICIs, for the elements detected in the two different bacteria datasets. **H)** Distribution of the pairwise wGRR values between all the genomes of PICIs. **I)** Symmetric heatmap of the matrix of the wGRR values ordered using hierarchical clustering. The colours follow the same code as in (**H**) with blue pixels representing low wGRR values (dissimilar genomes) and red pixels representing high wGRR values (similar genomes). The columns to the left of the heatmap indicate the bacterial species where the PICI genome was detected, the Type (A or B) of the PICI, and the component that is missing for given variants (exclusively for elements of Type B). The putative elements that were initially rejected by MacSyFinder, but that we included in the analysis, do not form a particular, segregated subcluster (**Fig S6**), suggesting they might be *bona fide* PICI.

PICI are present in many more species (190) than P4-like elements (81). These hosts are very diverse, including bacteria where phage satellites were not previously known to exist, and that are important in clinical and/or ecological settings. For instance, we detected putative PICI in *Acinetobacter, Bacillus* (as well as *Lactobacillus*), *Burkholderia, Clostridium, Rhodococcus* or *Sinorizhobium*. However, as most elements are found in a few (over-sequenced) bacterial genus (e.g., *Escherichia, Mycobacterium* and *Staphylococcus*), the difference in diversity considering the overrepresentation of PICI in certain bacterial taxa is less marked (Shannon Index = 2.55 for hosts of PICI elements versus 2.14 for hosts of P4-like elements). Some bacterial genomes can have up to five PICI, even if most bacteria have one or two of these elements (**Fig 2C**).

The organization of the PICI core components is well conserved, as suggested by previous studies and mentioned above (12) (**Fig 2E** and **Fig S4**). Only one of the most common variants shows a different order (Type B#var03, with a missing/unidentified primase-replicase), where the small terminase gene tends to be found before the capsid. We used the region from the integrase to the last identified core component to delimit the PICI (the element lacking the integrase was discarded). The resulting 1435 PICI have a median size of 9.5Kb (15 proteins), and only a few elements (17) have between 20 and 28Kb (**Fig 2F, Fig 2G**).

PICI are much more diverse, as measured by the wGRR, than P4-like satellites (p=0, two-sample Kolmogorov-Smirnov test). Their set of core components is smaller and most pairs of PICI have a very low wGRR (**Fig 2H**). The wGRR matrix groups PICI in four very distinct sub-families, each predominantly associated with a bacterial clade: *Escherichia* (two sub-families), *Mycobacterium*, and *Staphylococcus* (**Fig 2I**). Most elements in Mycobacteria tend to be very similar, forming two clusters with either an unidentified regulator or a primase-replicase. All the genomes of the putative satellites in *M. tuberculosis* (which are the majority of putative phage satellites identified in this taxon) are very similar to the previously described “small prophage-like elements” PhiRv1 and PhiRv2 (39, 40). Yet, other putative Mycobaterial phage-satellites are genetically distinct from the latter, and found across different species (e.g., in *M. abscessus*) (**Fig S5**). The *Staphylococcus* PICI lack a capsid modification gene, as previously described (12), and are split in many smaller subgroups. Those of *Escherichia* are very divergent and form smaller subgroups. There are also many small clusters of PICIs that each represent those few elements found in the genomes of other bacteria, and most of these PICI have four identified core components.

There are relatively few obvious cases of putative intra-species transfer of PICI. Only 73 pairs of elements with a high wGRR are found in different bacterial species (0.1% of the total number of pairs with wGRR > 0.9, n=60107). A large fraction is found in two *Staphylococcus* spp., although we do find some rare cases of putative transfer between more distant bacteria (e.g., *Acetivibrio* and *Tissierellia*). Thus, relative to P4-like satellites, PICI seem to be less frequently transferred across phylogenetically distant bacterial hosts. This may be a consequence of the presence of a majority of these elements in three very distantly related bacterial genera. Overall, these results uncover a plethora of very diverse PICI that are present in a large range of bacterial hosts.

### Capsid-forming Phage Inducible Chromosomal Islands (cfPICI) are a novel and distinct type of PICI

The cfPICI are a novel family of satellites related with PICI, but with a unique trait: they assemble their own cfPICI-specific capsid (8). Yet, cfPICI are incapable of forming viable phage particles because they lack other structural genes that they hijack from the helper phage, *e*.*g*. holins and tail-associated proteins. The presence of structural genes in cfPICI makes them more difficult to discriminate from prophages. For this reason, we defined profiles associated with phage tails, or phage holins, as forbidden components in the cfPICI model.

Five core components of cfPICI are homologous or analogous to the five core components of PICI, and in some cases the leftmost part of cfPICI (comprising the integrase, regulator and primase) were found to be exactly the same as PICI (8), suggesting they are indeed evolutionarily related. Some others are occasionally also found in PICI. This is the case of a nuclease (HNH) that is essential for phage head morphogenesis (and DNA packaging) in fully functional phages (41) and a head decoration module (a serine protease). We postulate that some of these genes might be used for the modification and stabilization of capsid morphology for phage capsids hijacked by PICI. However, there are several specific core components of cfPICI that allow to distinguish them from PICI: genes involved in the attachment of capsid to the hijacked tails (head-tail adaptor and head-tail connector), and a gene encoding the large terminase protein (*terL*) (**Fig S1C**).

We detected 969 cfPICI, all but three in chromosomes. Among these cfPICI, 177 had all the 11 core components (Type A), 459 had 10 (Type B), and 333 had 9 (Type C) (**Fig 3A and 3B**). 73 of cfPICI (<8%) were initially rejected by MacSyFinder due to the presence of a nearby gene homologous with either a single tail or holin gene. They were retrieved in the post-processing step because the forbidden component is outside the cfPICI (see Methods). Type A cfPICI were exclusively found in Proteobacteria. They are very abundant in Enterobacteriaceae (13% of all genomes), and in Pasteurellaceae (8%) and Morganellaceae (7%) (**Fig 3D**). Variants of cfPICI within Firmicutes tend to have the capsid component merged with the prohead serine protease component (and therefore genes matching exclusively the capsid can be absent). As MacSyFinder reports the best scoring profile for each gene, the latter is chosen instead of the capsid profile(s). These cfPICI are frequently found in Lactobaciiliaceae (25%) and Enterococcaceae (25%), and rarer (<2%) in Bacillaceae and Streptococcaceae. Overall, cfPICI are found in more species than PICI or P4-like elements (136, Shannon Index = 3.1), and are again present in several important species, where phage satellites have not been detected (e.g., from the genus of *Bacillus, Bordetella, Citrobacter, Haemophilus, Pseudomonas* or *Xanthomonas*).

**Fig. 3.**
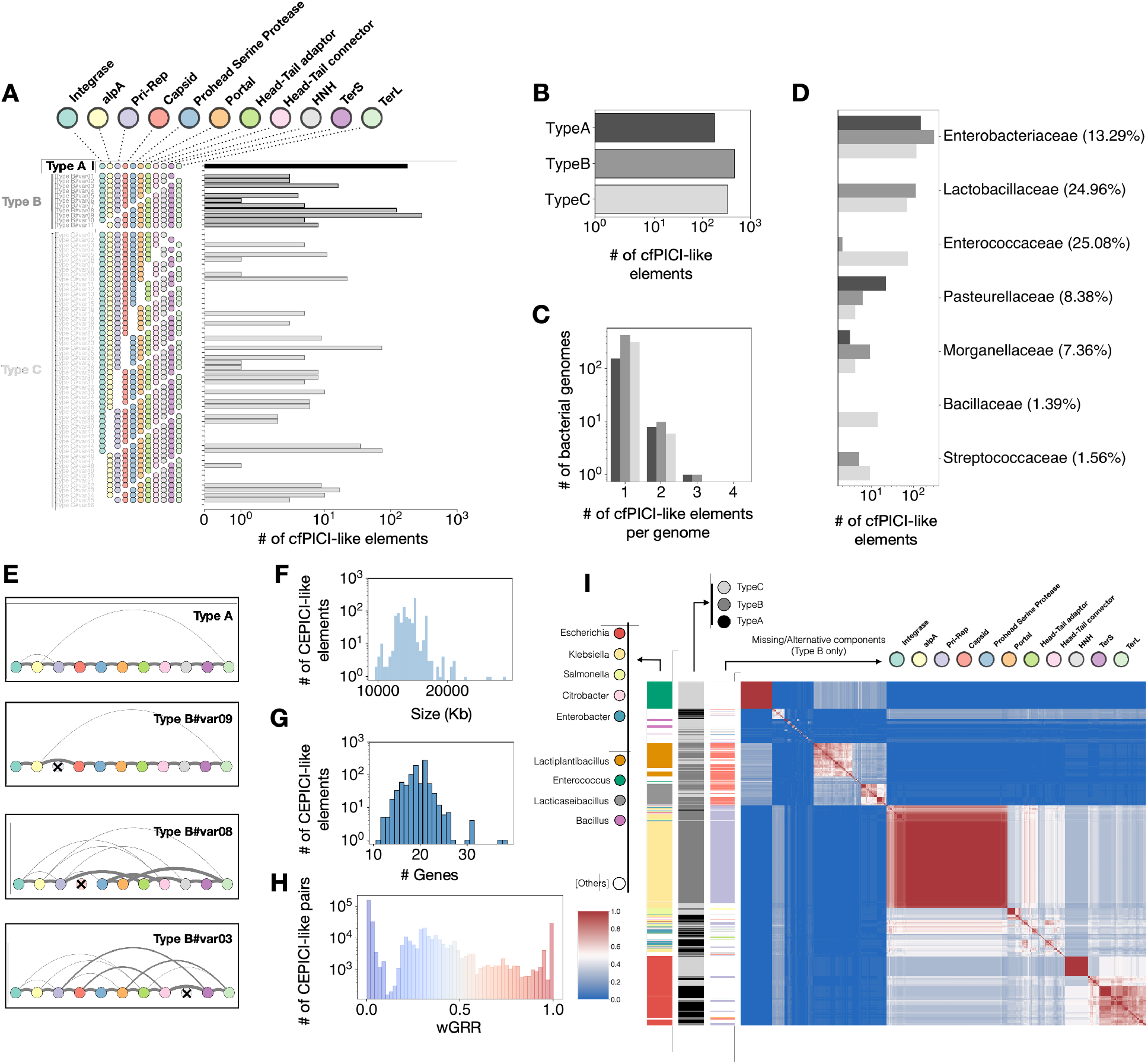
The abundance, genetic organization, bacterial hosts and genomic structure of the cfPICI family. **A)** Number of the different variants of cfPICIs identified in bacterial genomes. This includes elements that were initially rejected by MacSyFinder but recovered with our post-processing analysis, and correspond to 38, 15 and 20 of Types A, B and C, respectively) **B)** Total number of the different types of cfPICIs. **C)** Total number of cfPICIs per bacterial genome. **D)** Distribution of cfPICIs in bacterial families. The percentages in front of each family correspond to the proportion of genomes of that family where (at least) one cfPICI was inferred. **E)** Genomic organization of the four most frequent variants of cfPICIs. Nodes correspond to the different markers of the satellite (variants where the marker is absent have that indicated as a crossed-out node) and the edges between the nodes represent the frequency with which those two components are contiguous (but not necessarily adjacent). **F)** Distribution of the sizes of the extracted genomes of cfPICIs. **G)** Distribution of the number of proteins contained within each cfPICI genome. **H)** Distribution of the pairwise wGRR values between all the cfPICIs. **I)** Symmetric heatmap of the matrix of the wGRR values ordered using hierarchical clustering. The colours follow the same code as in (**H**) with blue pixels representing low wGRR values (dissimilar genomes) and red pixels representing high wGRR values (similar genomes). The columns to the left of the heatmap indicate the bacterial species where the cfPICI genome was detected, the Type (A, B or C) of the cfPICI, and the component that is missing for given variants (exclusively for elements of Type B).

The genomic organization of the core components of cfPICI is very conserved, with the exception of two variants that are more diverse (**Fig 3E** and **Fig S7**). We extracted the proteomes of the putative cfPICI, defined between the integrase and the farthest core component, after discarding five unusually large elements (>30kb) and the 48 elements without integrase. The remaining 916 putative cfPICI have a median size of 14Kb and encode a median of 19 proteins.

The gene repertoires of cfPICI are more similar than those of PICI, with a median wGRR of 0.2 and almost 10% of pairs with a wGRR higher than 0.8. Clustering the cfPICI by their wGRR reveals several subfamilies, usually associated with either monoderms (e.g., *Enterococcus* and *Lactobacillus*) or Proteobacteria (mostly *Escherichia, Klebsiella, Salmonella* and *Citrobacter*). This fits the previously obtained phylogeny of these elements (8). Within these sub-families, cfPICI tend to be more similar between closely related hosts. The cfPICIs added by the post-processing script (e.g., with neighboring prophage genes) integrate the existing clusters, suggesting that they are valid elements (**Fig S8**). Although there is a strong association between cfPICI subfamilies and particular bacterial hosts, c.a. 9% of the cfPICI pairs with a high wGRR (>0.9, n=36173) were detected in different host species. For instance, some cfPICI of *K. pneumoniae* are very similar to those found in *Salmonella, Enterobacter* or *Citrobacter*. This suggests that, relative to PICI, cfPICI are potentially capable of disseminate across more phylogenetically distant hosts.

### Phage-inducible chromosomal island-like elements (PLEs) are mostly clonal and are specific to *Vibrio cholerae*

PLEs were described in *Vibrio cholerae*, where they play a critical role in the defense from the virulent phage ICP1 (6). So far, all described PLEs are specific to *V. cholerae* (19) even if recently other putative satellites with homology to a few PLE genes were described in other Vibrio species (e.g., *Vibrio parahemolyticus*) (42). PLEs excise from the chromosome and package their genomes by hijacking ICP1. The cost for ICP1 is exacerbated by the acceleration of lysis promoted by PLEs after their packaging (43), which effectively halts the spread of ICP1 in the population.

To determine the core components of PLEs, we first selected the homologs present in at least three of the five prototypical PLE genomes (PLE 1 to 5 (6)). We treated differently the components most distant from the integrase because they are more variable; hence, we selected those found in at least two of the five prototypical PLEs. We used the core components for the iterative search of putative PLEs in the Vibrionaceae genomes (see Methods), which resulted in the selection of a large number (15) of highly frequent markers for PLE. Some of these markers have a well-defined role in the PLE lifecycle: an integrase, a gene that represses the capsid morphogenesis of ICP1 (*capR*), a replication initiation protein (*repA*), a nickase that hampers the replication of the hijacked phage (*nixI*), and a gene that accelerates the lysis of the bacterial host cell (*lidI*). Other highly frequent markers of PLE to which we were able to assign a functional PFAM annotation include a protein with an HTH binding domain, which was previously described in PLEs (6); a sigma 70-like factor, a component of the specificity subunit of the bacterial RNA polymerase; and a profile with homology to a cyclin-dependent kinase-activating kinase (MAT1) suggested to be involved in nucleotide excision repair of damaged DNA (44). Seven other highly frequent markers (M1 to M7) were uncharacterized, and we were unable to annotate them using PFAM. Given the substantial variation in terms of presence/absence of these markers in the known PLE genomes (see **Table S1**), we used all the 15 markers to study the natural variation of this satellite family.

PLE are specific of a few Vibrio and are rare in our original dataset. To increase the sample size, we retrieved from Genbank all complete and draft genomes of Vibrionacea (11627 genomes, see Methods). We detected 410 elements of Types A to I, i.e., with between all (Type A) and 7 (Type I) PLE markers (**Fig 4A**). Most genomes have a single PLE. Some of these types correspond to known variants of previously identified PLEs. Elements of Type A (n=238) correspond to PLE1 and elements of Type B (n=38), for which we only find a single variant, correspond to PLE5. PLE2 and PLE4 correspond to two different variants of Type D and PLE 3 corresponds to a variant of Type F (**Table S1**). The more incomplete elements (Types G, H and I) tend to either lack or have highly distinct first or final half of the PLE markers we assembled, and it is likely that they result from assembling artifacts inherent to draft genomes. All the putative PLE-like satellites detected in the Vibrionaceae dataset were found in *V. cholerae* (in 12% out of its 3446 total genomes for this species).

**Fig. 4.**
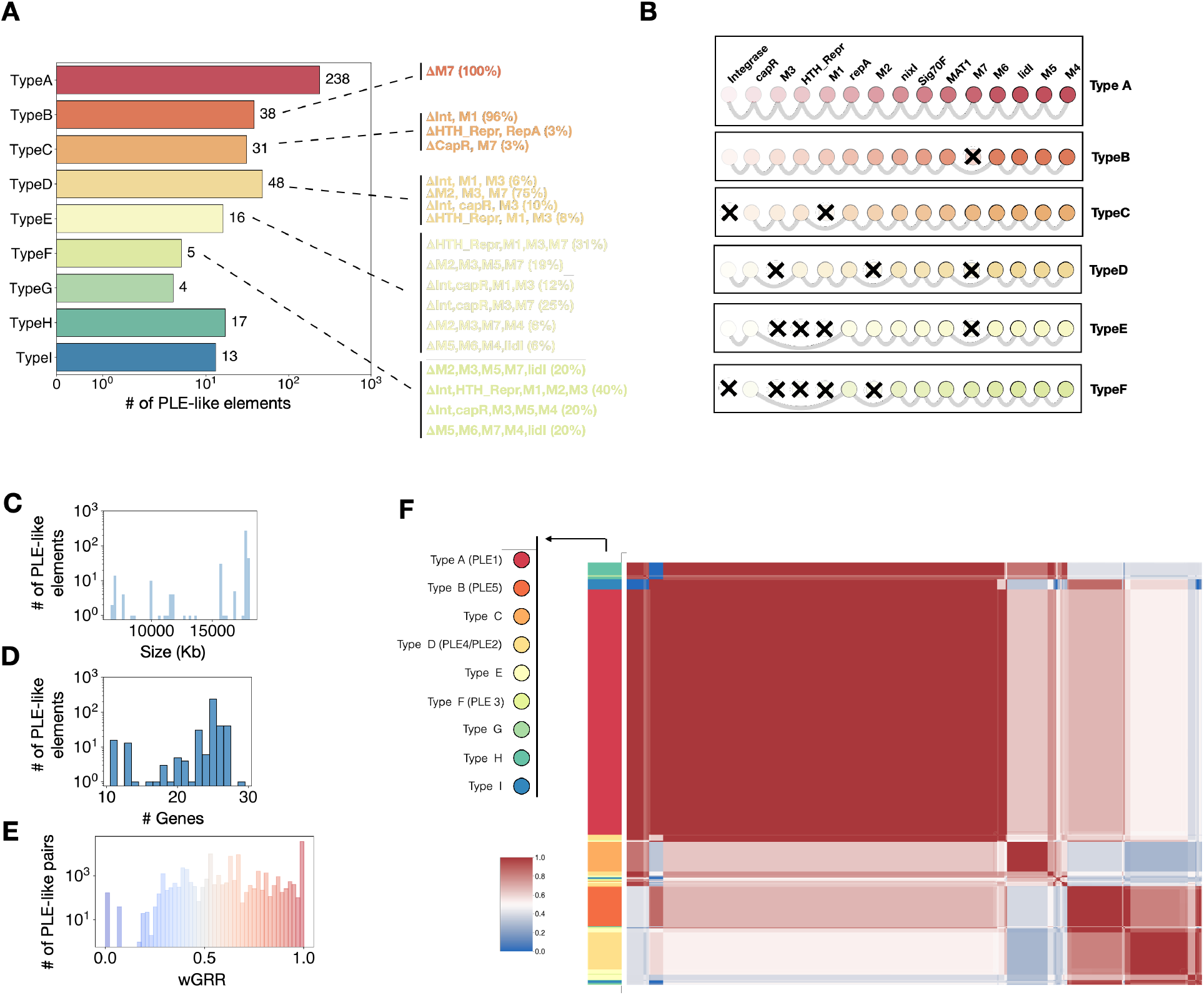
The abundance, genetic organization, genomic structure of PLEs. **A)** Number of the different types of PLEs identified in Vibrionaceae genomes. Dashed lines in front of the bars indicate the variants within each type, more specifically which components are missing/undetected. The proportion of each variant is shown in parentheses. **B)** Genomic organization of the most frequent variant for each PLE type. Nodes correspond to the different markers of the satellite (variants where the marker is absent/unidentified have that indicated as a crossed-out node) and the edges between the nodes represent the frequency with which those two components are contiguous (but not necessarily adjacent). **C)** Distribution of the sizes of the extracted genomes of PLEs. Although we extracted genomes for all identified sets, we do not use those that are found between contigs (1.2%) to account for the distribution in genome size, as the precise genomic locations (and relative distances) of the proteins would be unreliable. **D)** Distribution of the number of proteins contained within each PLE genome. **E)** Distribution of the pairwise wGRR values between all the PLE genomes (in both bacterial datasets). **F)** Symmetric heatmap of the matrix of the wGRR values ordered using hierarchical clustering. The colours follow the same code as in (**E**) with blue pixels representing low wGRR values (dissimilar genomes) and red pixels representing high wGRR values (similar genomes). The column to the left of the heatmap indicate the type of PLE (in parenthesis, the prototypical PLEs that are classified with a similar type).

The core components present across the PLE variants are very diverse (**Fig 4B**), but the order of the components is extremely well conserved (**Fig 4C**). We extracted the region between the integrase and the furthest component (typically M4) in Type A elements. These regions have a median size of 17.7Kb (and 25 proteins). These elements tend to be genetically very similar, with more than 90% of the pairs having wGRR values higher than 0.9. Further, 12 genomes are enough to regroup all other satellites at wGRR≥0.9, i.e., the remaining genomes are very similar to one of these (**Fig 4F**). While this may seem contradictory with the observation that several core genes are often missing, PLE differ from the other elements in that a large fraction of genes are core. Some of these 12 groups were much more abundant than others, leading to a few very large clusters (**Fig 4F**). Overall, our results further confirm that PLEs are very distinctive from other satellites and have limited genetic diversity, forming highly related sub-families around the previously known PLE types.

### Phage-satellite families form genetically distinct groups of mobile elements and can co-exist in bacterial genomes

All satellites exploit functional phages for mobilization, but the mechanisms involved in this process differ widely. This raises questions regarding the evolutionary origin of these elements, in particular whether they diversified from a single ancient satellite, or if they evolved multiple independent times. To understand the similarities between the different families of satellites, we analysed the co-occurrence of core genes from all satellite families across all the putative phage satellite genomes identified. For each satellite, we note the presence/absence of each core gene. The resulting binary matrix with all satellites was clustered (**Fig 5**) and revealed that three of the four families (P4-like, cfPICI and PLE), form well-separated and cohesive clusters of elements. PICI aggregated in multiple clusters, which is consistent with their high gene repertoire diversity (**Fig 2I**). Interestingly, the separation of satellites in different clusters is due to the combination of their markers, and not due to the presence or absence of a single one. For instance, AlpA is found in all P4-like, in many cfPICI and in some PICI. Moreover, the same gene can match two regulatory profiles (e.g., *alpA* and *merR*), although sometimes a single satellite genome can also have two regulatory genes, each matching a different protein profile. This analysis also occasionally identified satellites of a given family encoding components that are core from another. For instance, some cfPICI have a homolog of ε from P4, some P4-like satellites have a homolog of HNH from cfPICI, and some PLEs have a TerS homologous to that of PICI and cfPICI. These results show that the different families of satellites are clearly distinct, but they also suggest gene flow of core genes between satellite families.

**Fig. 5.**
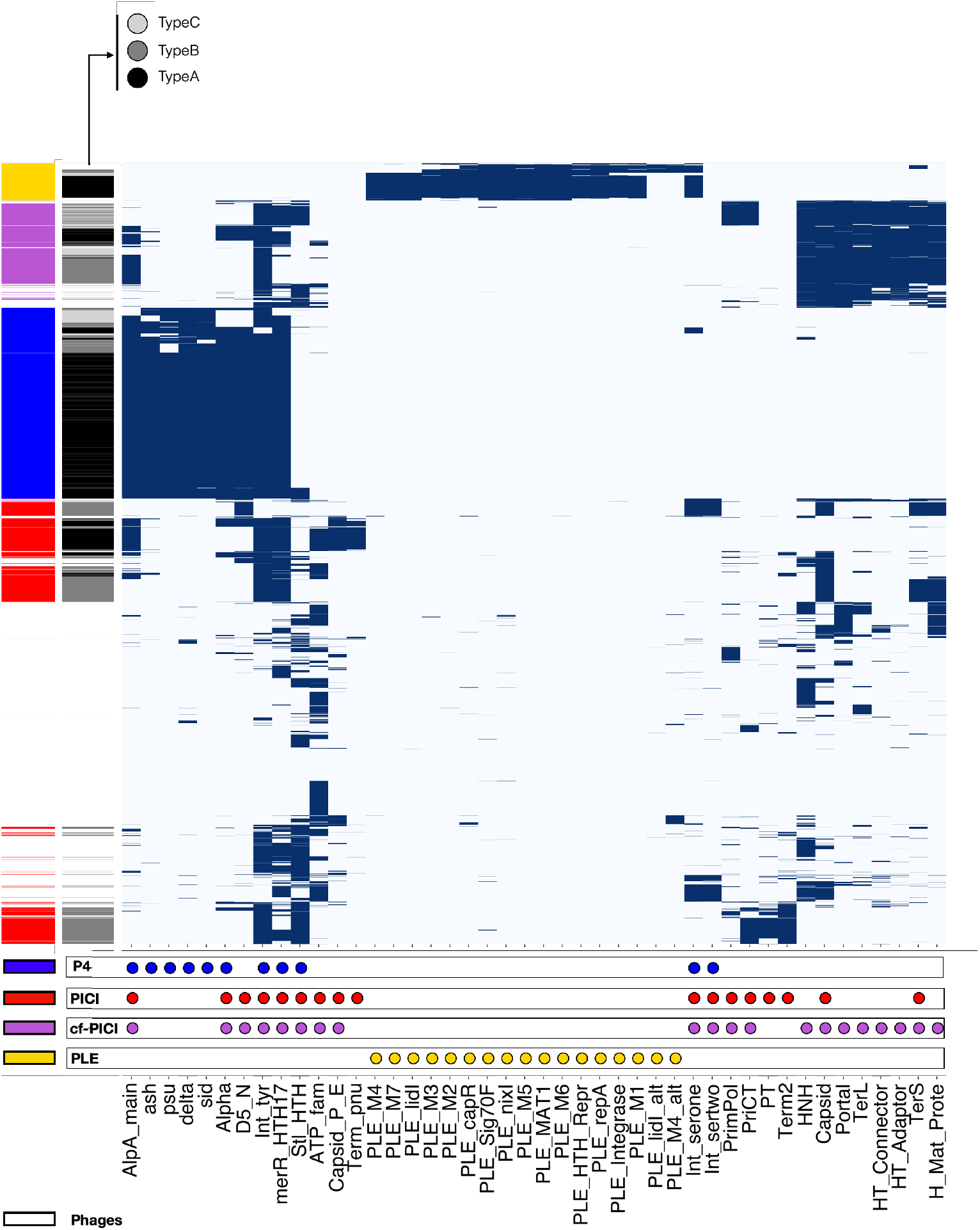
Pattern of presence/absence of the different satellite markers across all phage and putative phage-satellite genomes. Hierarchical clustering of each phage or phage-satellite genome (row) assessed for the presence of a given phage satellite marker (column, at least one homolog gene with an e-value of at most 1e^-2^ and minimum coverage of 40%). For each genome, if a homolog for a marker is present, the cell is shown as blue, otherwise its absence is shown as an empty cell. The markers used by SatelliteFinder to detect each phage satellite family are shown below the matrix as circles. The columns on the right of the matrix indicate (from right to left): the family of the phage-satellite genome corresponding to each row; and the Type of each satellite genome (limited to Types A, B and C).

Our analysis could mistakenly annotate phages as satellites, because they have homologous components. To test this, we retrieved the genomes of 3725 phages and made an analysis similar to that done with the satellites. Almost all phages have at least one homolog to a satellite core gene. This is expected because both often have an integrase and capsid-associated genes. Nevertheless, there are rarely more than a few of these core genes in common between phages and satellites. The clustering of the large binary matrix of presence of core genes in satellites and phages shows that the latter form clusters well separated from those of satellites (**Fig 5**). This confirms that our method discriminates the two types of elements, and that satellites are very distinct from phages.

We assessed the evolutionary relationships between PICI and cfPICI. In this particular case, the two families of satellites share several core components. We tested if SatelliteFinder is able to accurately distinguish them. First, no elements were simultaneously identified by the cfPICI and PICI models. Second, no PICI satellites were identified when searching for them directly in the dataset of cfPICI genomes. Finally, no cfPICI was identified when searching for them directly in the PICI genomes. This suggests we can discriminate them accurately. To confirm this, we quantified the genome-wide similarities between PICI and cfPICI, by computing the wGRR between them. The clustering of the wGRR values revealed little to no mixing between the major clusters of these two types of satellites (**Fig S9**). We then built phylogenetic trees for four homologous core components: the transcriptional regulator, the primase-replicase gene, the capsid gene (or capsid-modification gene in PICI) and the small terminase. This revealed contrasting results. On one hand, the trees of the transcriptional regulators and of the primase-replicase are strongly paraphyletic (**Fig 6A and Fig S10)**, suggesting gene flow between the two families of satellites (or intermingled evolutionary history). On the other hand, the capsid (**Fig 6B** and small terminase genes (**Fig S11)** separate almost perfectly PICI from cfPICI. This suggests that some functions are more likely to be exchanged between different types of satellites. Alternatively, the integrase-proximal genes may constitute a functional module that might be involved in the cross-regulation of (potentially similar) helper phages. These results confirm that the first half of the PICI and cfPICI satellites are often similar and suggest a common evolutionary history. Hence, these are two evolutionary related, but different, families of satellites whose part of the hijacking modules (those that *de facto* distinguish PICI from cfPICI) have evolved independently for a long time.

**Fig. 6.**
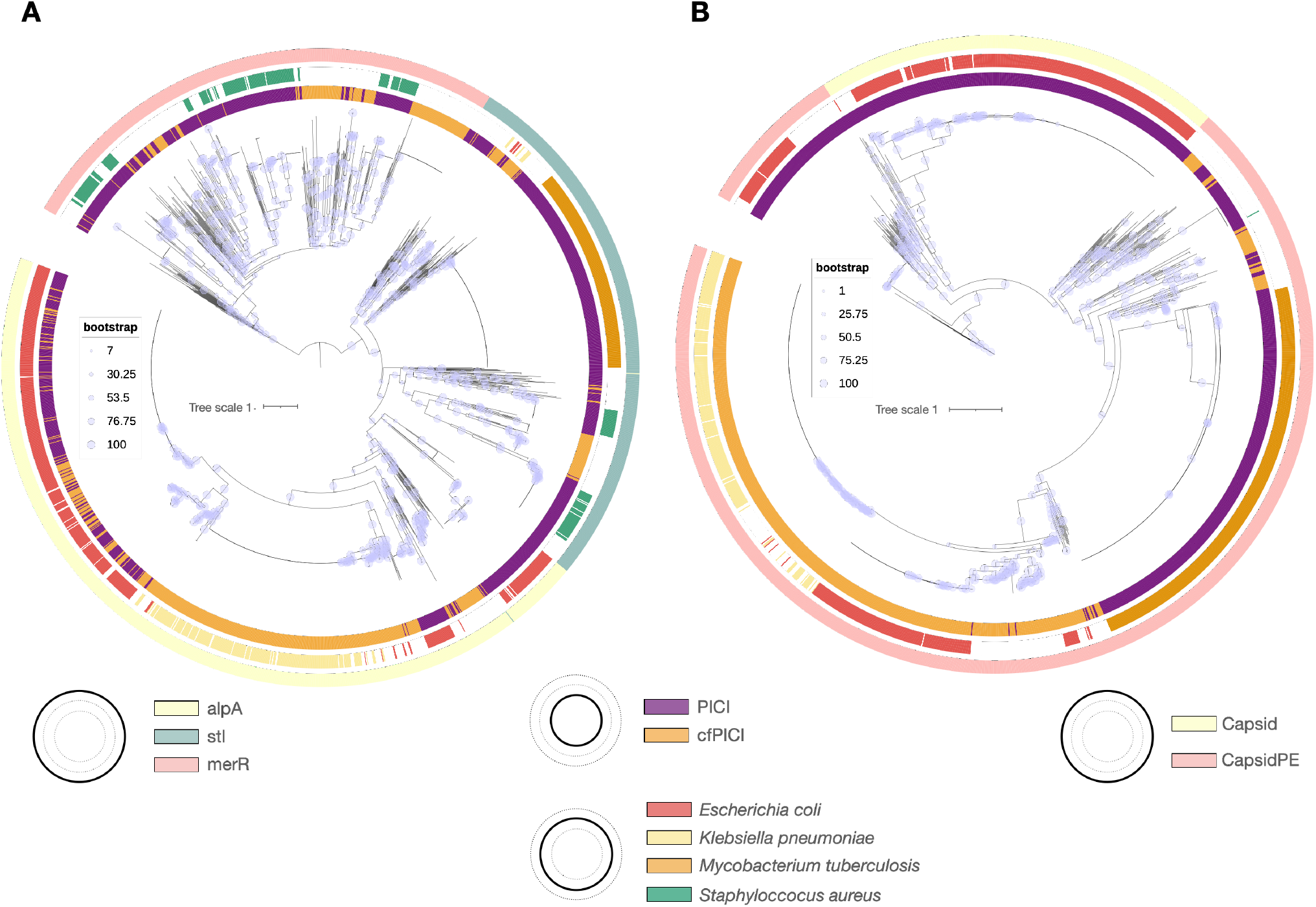
Phylogenetic trees of two of the prototypical components of PICI and cfPICI. **A)** Tree of the regulators of cfPICI and PICI. **B)** Tree of the capsid, or capsid-like components of cfPICI and PICI. In both A and B, outer circles indicate (from inside out) the family of satellites (PICI in purple, cfPICI in orange); the most frequent bacterial host species of the phage-satellites (less frequent host species are shown as white spaces); and the profile associated with the specific component (AlpA, MerR or StI for the regulator in A; in B, the two profiles used to detect capsid-associated components (capsid or capsidPE, see **Fig S1**)).

Finally, since the families of satellites can be separated accurately, we assessed if multiple phage satellites, each belonging to a different family, could co-integrate the same bacterial genome, or if they mostly exclude each other. PLE never co-occur with other satellites, which is justified by its narrow bacterial host range. However, we found that a few hundred bacterial genomes have at least one P4-like and one PICI (227 genomes) or a P4-like and a cfPICI (379 genomes). The former combination is mostly found in *E. coli* or *Shigella boydii* genomes, while P4-like and cfPICI satellites tend to co-integrate a more diverse set of hosts, from *E. coli* to *K. pneumoniae, Citrobacter freundii* or *Yersinia enterolitica*. Combinations of PICI and cfPICI are less common, and we found them in only 47 bacterial genomes. Impressively, 31 *E. coli* genomes have at least one element of each of the three families of satellites (P4-like, PICI and cfPICI). These results show that satellites of different families often co-exist within single bacterial genomes.

## Discussion

In spite of the impact of satellites in phage reproduction and in the transfer of adaptive traits among bacteria, there was little information in the literature on their composition and number. We were able to leverage previous curated analyses of a few dozens of such elements to build a tool – SatelliteFinder – that systematically and reproducibly identifies phage satellites in bacterial genomes. This approach revealed numerous novel satellites, highlighting their relevance in nature. Nevertheless, it has some limitations. First, the existence of few examples of experimentally validated satellites means that it is not possible to accurately evaluate our classification. We overcame this by using an iterative cycle of enrichment and curation of satellite markers, but this also means that the phage-satellite models we propose must be regarded as a first educated attempt to characterize each family. Interested users can easily modify MacSyFinder models by changing one single text file. This will facilitate the search for novel variants of known satellites and to add novel types of satellites. Second, the presence of mobile genetic elements in the close vicinity of phage satellites can affect the identification of the latter. For instance, multiple contiguous phage-satellites could be identified as an overly large single satellite and a neighboring phage may lead to the exclusion of a valid satellite because of more than a single nearby phage-like marker. The post-processing analysis we perform using the output of MacSyFinder is important in diminishing these issues, but does not always solve it. For instance, while this manuscript was being submitted, a preprint showed that satellites can integrate inside prophages (45), which poses an even more difficult scenario to accurately distinguish these elements from prophages. Future work will be needed to assess the ability of our approach to correctly assign such cases. Third, we assume that satellites are delimited between the two furthest (identified) components. This may result in an underestimation of the size of the satellite, as well as their genomic repertoires. For instance, some accessory genes in PICI, namely genes involved in anti-phage defense, are found between the attachment site and the integrase (10). Fourth, we cannot ascertain if all satellites are functional. One would expect that most elements with the full set of core genes are functional. This is less established for variants missing core genes. Yet, some of these variants are highly conserved and a few were experimentally validated, suggesting that many of them might be functional. Fifth, our approach is sensitive to the mis-annotation of small genes as pseudogenes. For instance, in our previous analysis, we inferred that a large number of Type B P4-like satellites were alpA-less, as most of them had an inactivated (pseudogenized) *alpA* gene. However, this variant is very rare in the current analysis, because the most recent bacterial database annotates *alpA* correctly. Finally, the PLE analysis was done with draft genomes. The identification of phage satellites from draft genomes, where they might be split across different contigs, or within misassembled contigs, can erroneously suggest that some elements are either missing or have alternative core components. This seems to be the case for the less complete PLE variants (Types H and I) that had homology to either the first or the last “half” of the PLE model. This is not an issue when using complete genomes, as was the case for the analysis of P4-like, PICI and cfPICI.

Both the commonalities and uniqueness of the different families of phage satellites provide insights on their function and evolution. Integrases are core genes of all satellites and almost all elements were integrated in the chromosome. Regulatory components are also present in almost all satellites, and they are much more satellite-specific than integrases. It has been suggested that AlpA (or its functional analogues) mediate the cross-regulation between satellites and helper phages (12, 16). The potential gene flow suggested by the mixed phylogenies indicates that there might be functional compatibility, both within and across satellite families (i.e. one regulator can be exchanged by another). It also raises the possibility that different satellites could exploit the same phages, independently of the satellites’ hijacking strategy. Hence, regulators might be one of the best candidates to understand the evolution of satellites as a large family of elements. Another candidate could be the primase-replicase component(s), which are present in all of the families studied here, but the divergence between them suggests that they are analogs instead of homologs. Other components seem to be more specific to each family of phage satellites. The Psu-Delta-Sid operon of P4-like phage satellites performs a unique and conserved helper subversion strategy; the capsid assembly and head-tail adaptor genes of cfPICI means that this is the only satellite known so far that hijacks only part of the helpers’ virions; and PLEs have evolved genetic machinery to not only subvert a specific incoming virulent phage, but also to kill its bacterial host upon excision. Interestingly, this strategy might also be used by some PICI, that were recently described to encode an abortive infection system (10), and P4-like satellites can also encode (less drastic) anti-phage defense systems (e.g. retrons) (9). Hence, while the overall strategies (of subversion, replication or anti-phage defenses) of the different families of satellites might seem similar, they occur through different genes and mechanisms, and are likely to have evolved multiple times.

Novel, and experimentally testable insights into the different core functions of phage satellites can also come from the presence/absence and organization of their core components. The conserved genetic organization of most satellites, including its variants, suggests the existence of tight relationships between contiguous core components and conserved programs of gene expression. Certain variants lack core components and form specific clades, thus suggesting they might be functional. The latter would be an indication that these components are facultative. For instance, PICI of Type A represent a sub-family that relies on capsid modifications to hijack their helpers, while those of Type B do not require such a function. Psu-less variants of P4-like satellites are also frequent and form sub-families. One of these is associated with a specific bacterial clade (Serratia), suggesting that its function is either unnecessary or complemented by other components (e.g., Sid), and might have evolved within the bacterial species and in the context of its prophages. Other variants are rare and integrated into existing subfamilies of more complete types. These cases could represent recent loss of core components, that result in defective variants and hence truly essential components of these satellites.

The different families of satellites are strikingly different in their abundance, taxonomic distribution, genetic composition, and genetic diversity. All of these might be suggestive of the ecological conditions that underlie the establishment of the tripartite relationships between bacteria, phages and their satellites. For instance, the reduced diversity and narrow bacterial taxonomical range of PLEs might result from the high conservation of ICP1 making it unlikely to infect (and thus transfer PLE to) other bacterial species (46). However, since PLEs were not described to exploit other phages, virulent or lysogenic, this also suggests that the tremendous selective pressure of ICP1 on *V. cholerae* might have created the conditions for the tight and highly specialized function of PLEs in the ecology and evolution of this bacterial pathogen. Other satellite families show a much higher genetic diversity. P4-like satellites are found across many Enterobacteria, and PICI and cfPICI are particularly diverse and they have the hosts with broadest taxonomic span. It remains to be uncovered whether this results from a promiscuous relationship between these satellites and their broad host-ranged helper phages (as our data suggests it is the case for some P4-like satellites), or from the diversification of ancient versions of these satellites that diversified within each bacterial clade. Our results also show that many bacterial genomes have multiple satellites of the same family. This creates an interesting context for interactions between these elements. One recently described example of satellite-satellite interactions involves a specific PICI (SaPI3) that is induced not by prophages, but instead by other co-integrated PICI (47). Furthermore, individual bacterial genomes often carry satellites of different families, suggesting that competitive or antagonistic interactions between phage satellites of different families may occur. Together with the ubiquity of prophages, and the oft-present anti-phage defense systems in phage satellites (6, 9, 10), this highlights the complex networks that dictate the emergence and maintenance of these phage satellites and phages, as well as the fitness and survival of their bacterial hosts. Our results reinforce the idea that phage satellites play an important role in the microbial world. Even if our approach was conservative, requiring the presence of many core genes in common to known satellites, we detected ca 5000 phage-satellites in bacterial genomes. This number is huge, given the little we know about the distribution and diversity of these elements. There are even more elements that we excluded because they lacked too many core genes relative to the known satellites (e.g., we found more than 6000 Type C PICI elements). These elements contain several hallmarks of phage-satellites, *e*.*g*. AlpA or packaging proteins, which suggests there may be many other, still undescribed, families of satellite in bacterial genomes. Such elements may use novel exciting mechanisms for replication, sensing, or phage hijacking. Many elements in marine bacteria have recently been proposed to be phage satellites (17), and other some “incomplete” PLEs were recently described in Vibrionacea other than *V. cholerae* (42). Considering that half of the bacteria have at least one prophage (48) and that we identified satellites in a much smaller number of bacterial genomes, it is very likely that this is the beginning of the characterization of a vast diversity of phage satellites. SatelliteFinder allows to easily add or modify models for satellites. It allows to experiment combinations of both known and hypothetical marker genes, which will be key to identify novel putative satellites for experimental verification.

## Acknowledgements

We thank Kim Seed for comments and suggestions on earlier versions of the manuscript, Graham Hatfull for helpful discussion regarding the PhiRv1 and PhiRv2 elements in *M. tuberculosis*, and Bertrand Néron and Fabien Mareuil from the Institut Pasteur’s Bioinformatics and Biostatistics Hub for help in the development of the Docker and Galaxy version of SatelliteFinder. We acknowledge funding from Equipe FRM (Fondation pour la Recherche Médicale): EQU201903007835, Laboratoire d’Excellence IBEID Integrative Biology of Emerging Infectious Diseases [ANR LBX-62 IBEID AAP BOURSE S2I ROCHA]. This work used the computational and storage services (TARS cluster) provided by the IT department at Institut Pasteur, Paris.

## Data availability

The bacterial and phage genomes, as well as most profiles used to detect the core components of phage satellites, are publicly available. For the core components without public HMM profiles, we include the custom profiles as Files S2 to S19. The models used for MacSyFinder are also available MacSyModels in the public repository. The additional custom Python scripts to post-process the output of MacSyFinder are included in the Docker image at (https://hub.docker.com/r/gempasteur/satellite_finder).

