## SupplementaryFigures for "Identification and characterization of thousands of bacteriophage satellites across bacteria"

**This PDF file includes:**

Figures S1 to S11
Table S1

**Other supplementary materials for this manuscript include the following:**

Files S1 to S19

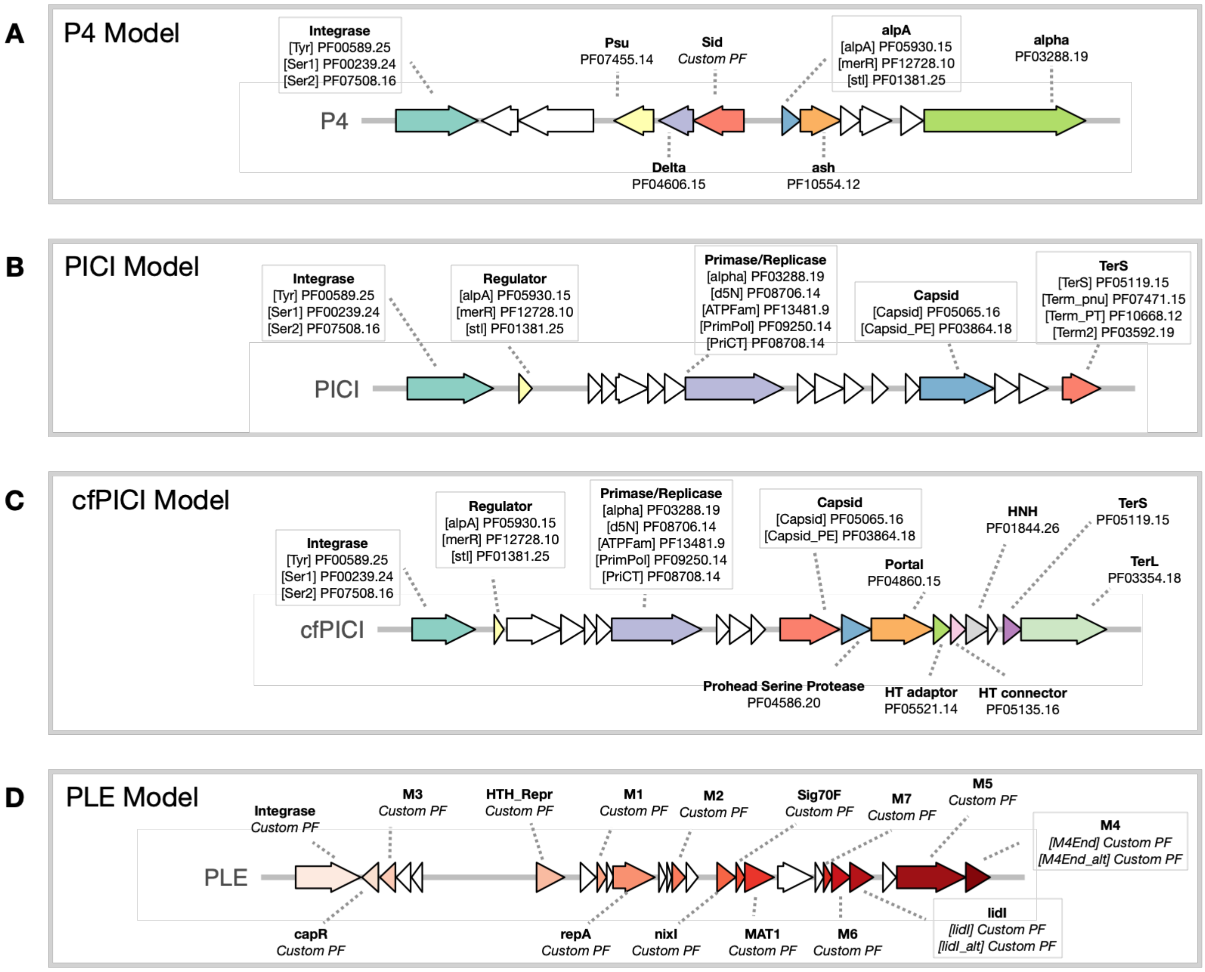

**Fig S1. Models for each of the satellite families**

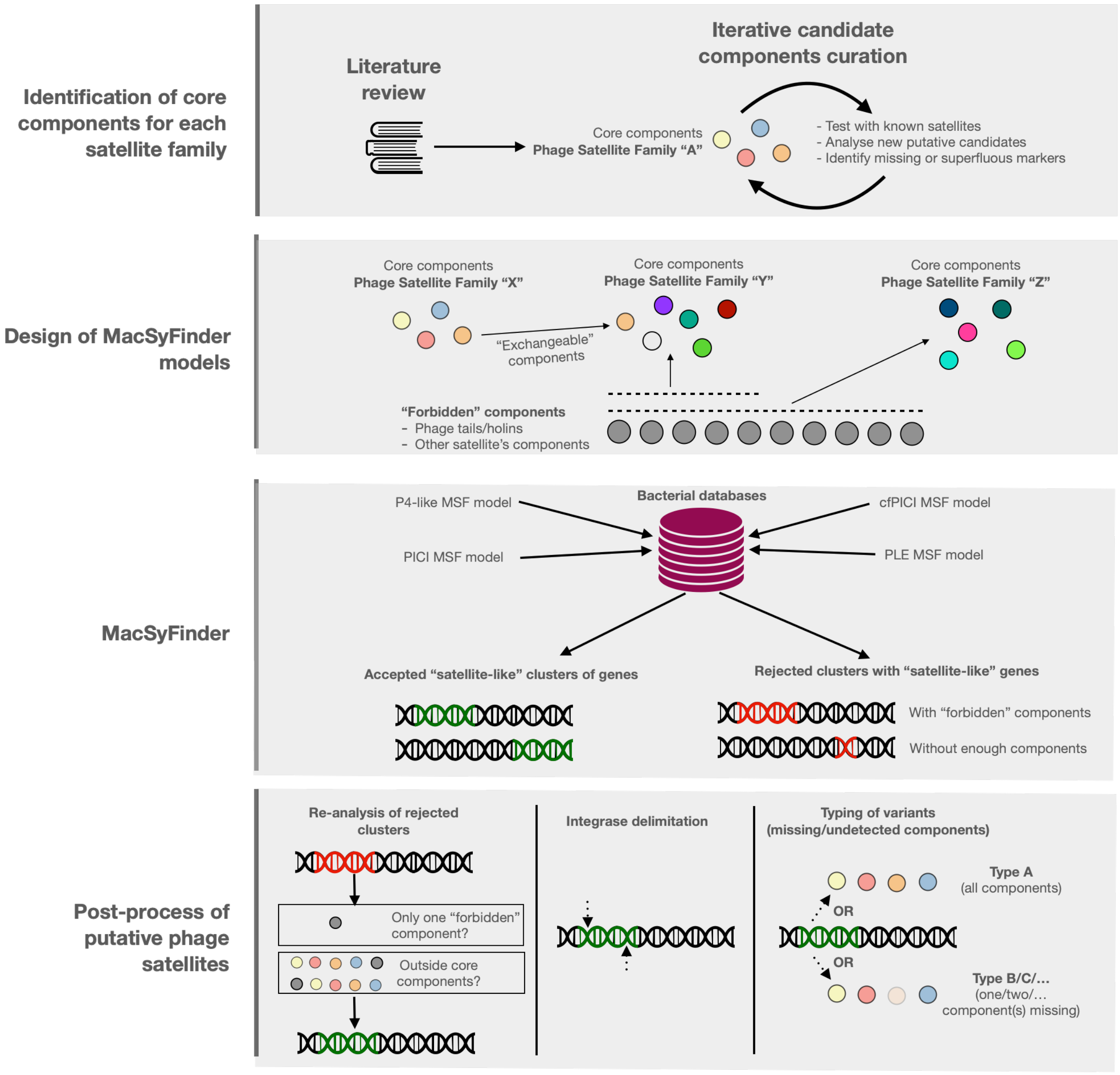

**Fig S2. Summary of overall process**

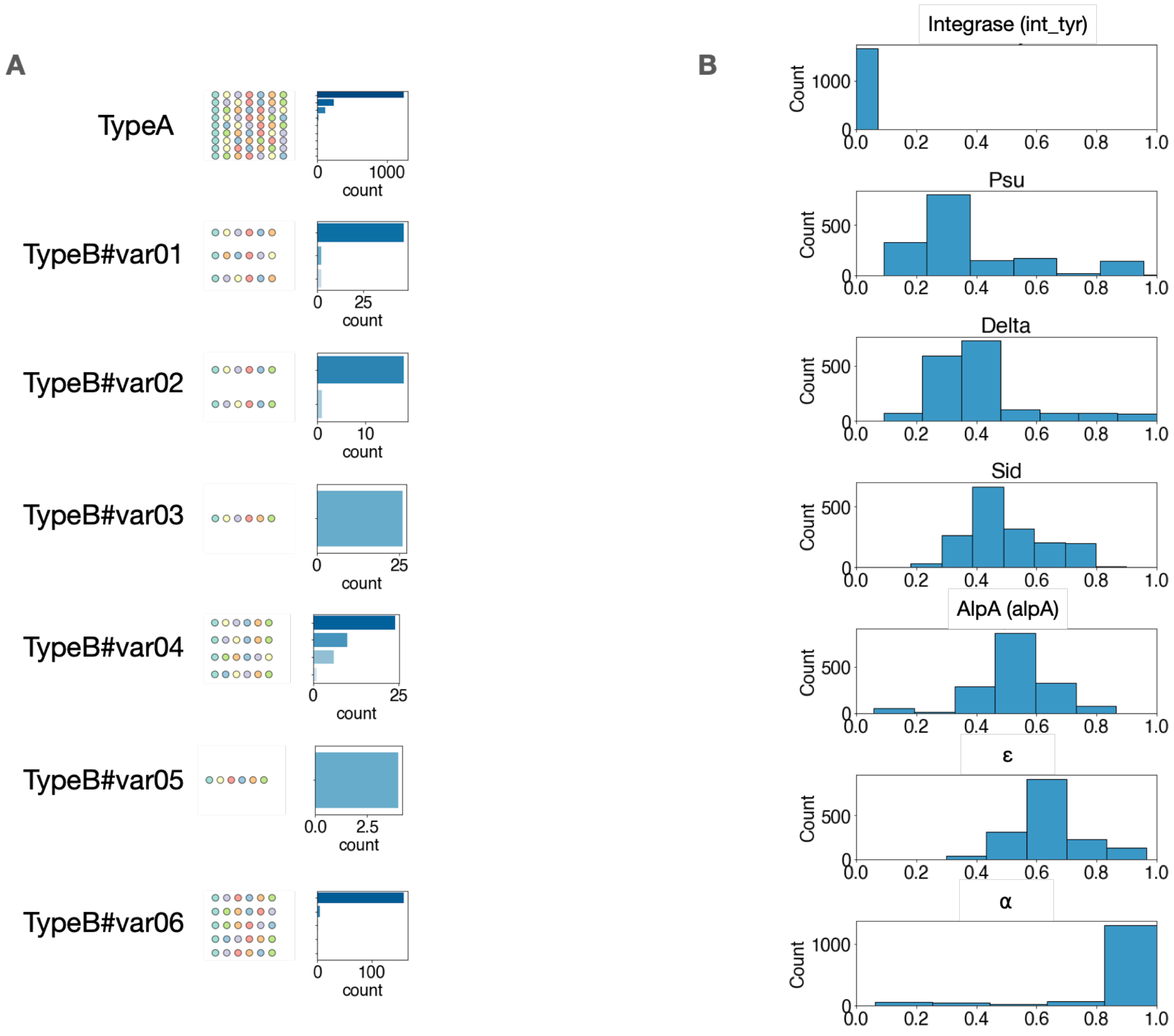

**Fig S3. Detailed genomic organization of P4-like satellites**

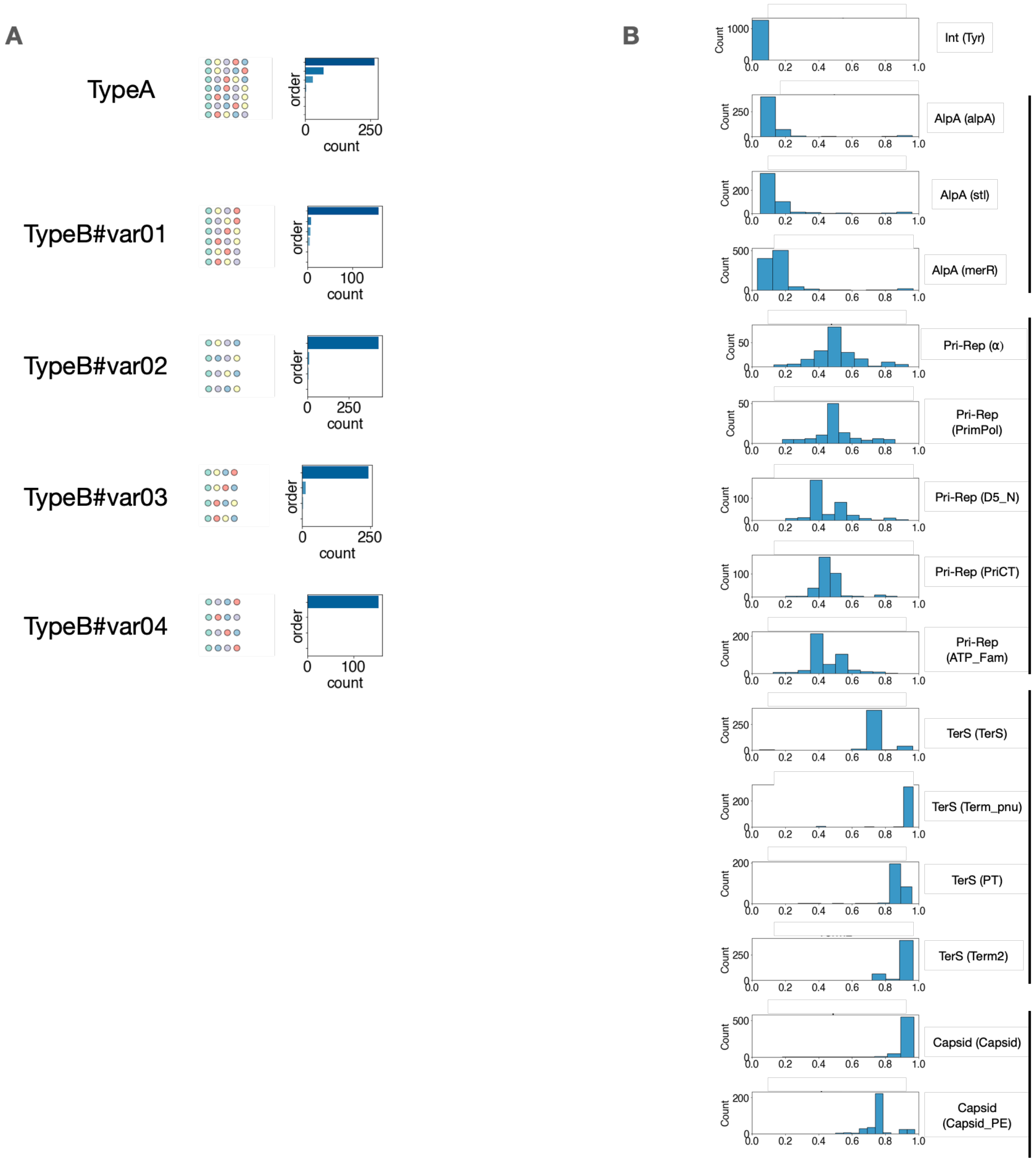

**Fig S4. Detailed genomic organization of PICI**

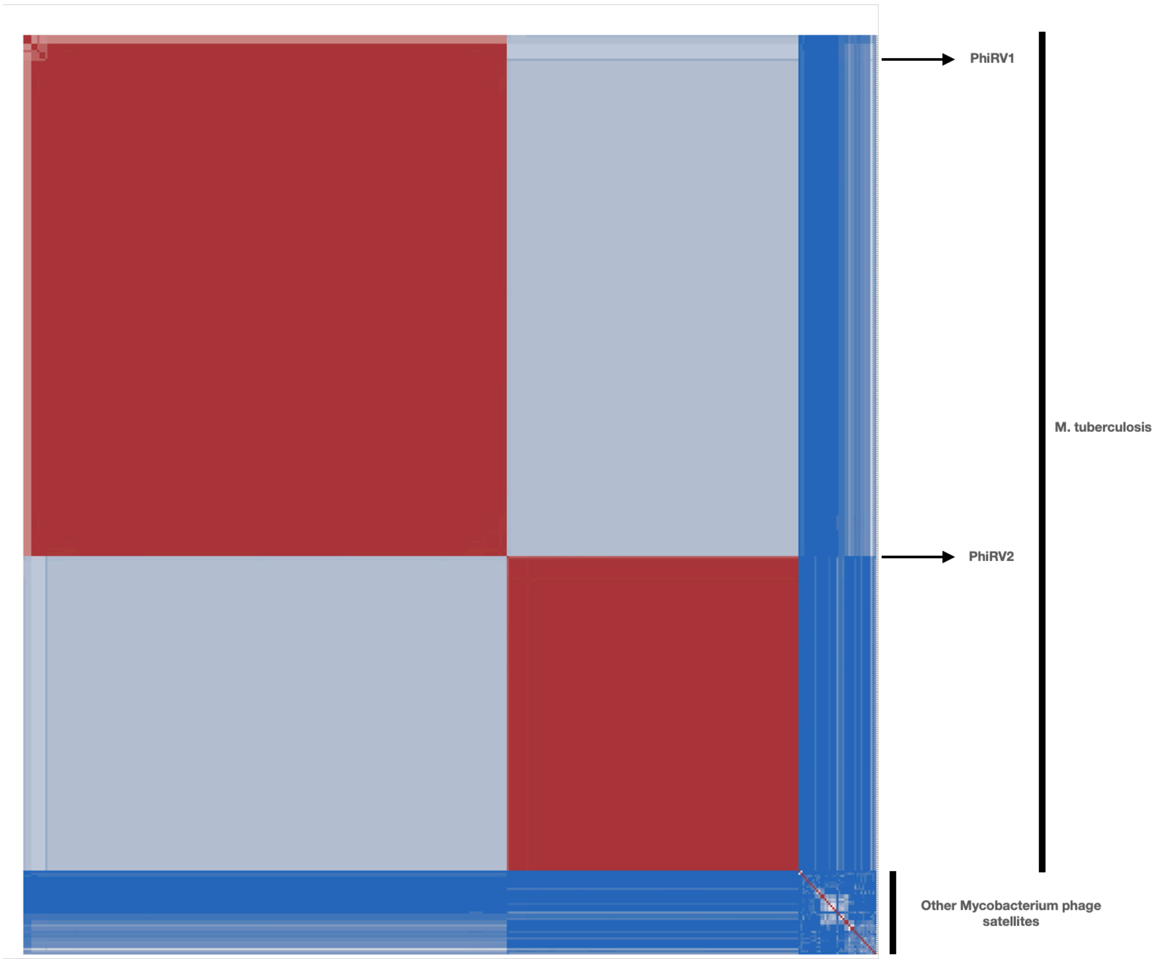

### Fig S5. wGRR comparison of Mycobacterial PICI

Symmetric heatmap of the matrix of the wGRR (weighted gene repertoire relatedness) values between pairs of PICI identified in Mycobacterial species (as well as PhiRv1 and PhiRv2), ordered using hierarchical clustering. Blue pixels representing low wGRR values (dissimilar genomes) and red pixels representing high wGRR values (similar genomes).

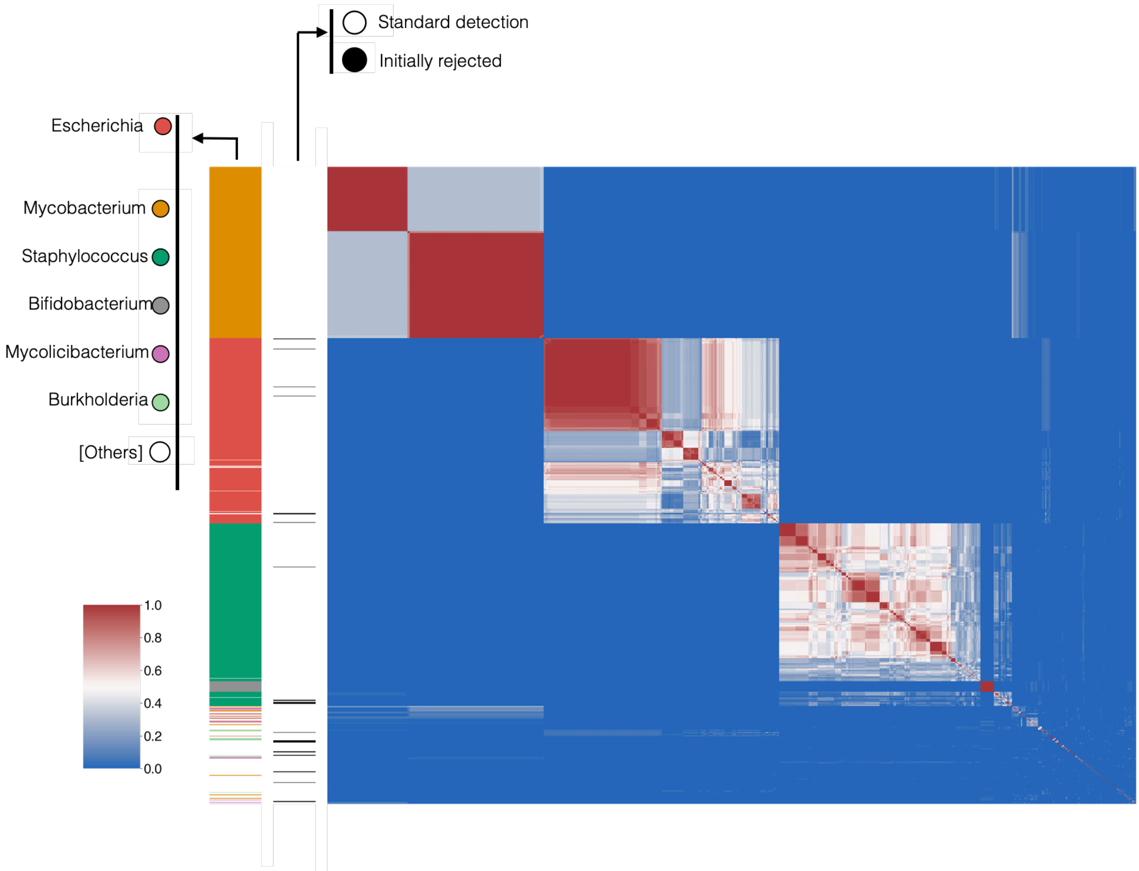

**Fig S6. Localization of initially rejected PICIs within the wGRR clusters of the family's genomes**

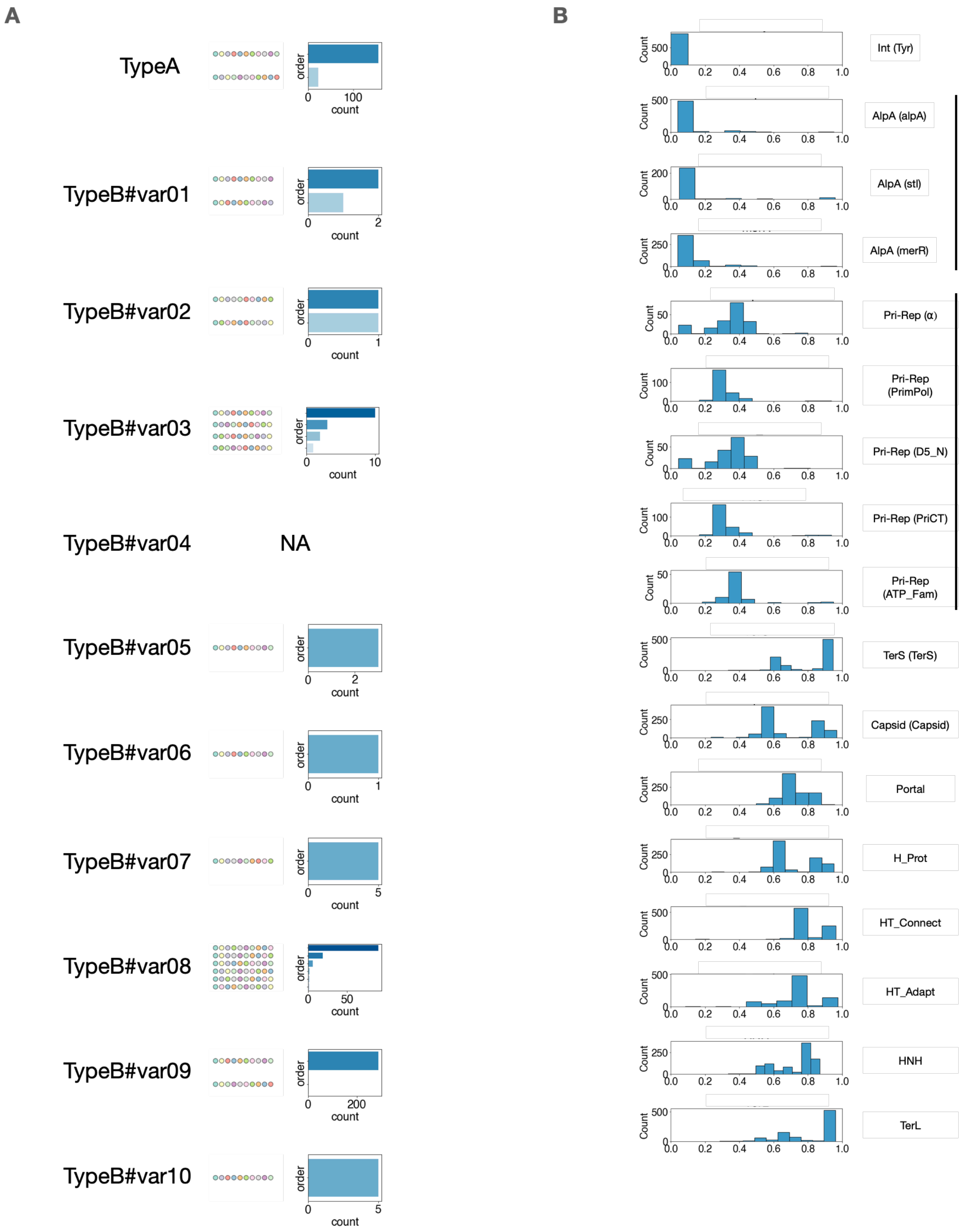

**Fig S7. Detailed genomic organization of cfPICI**

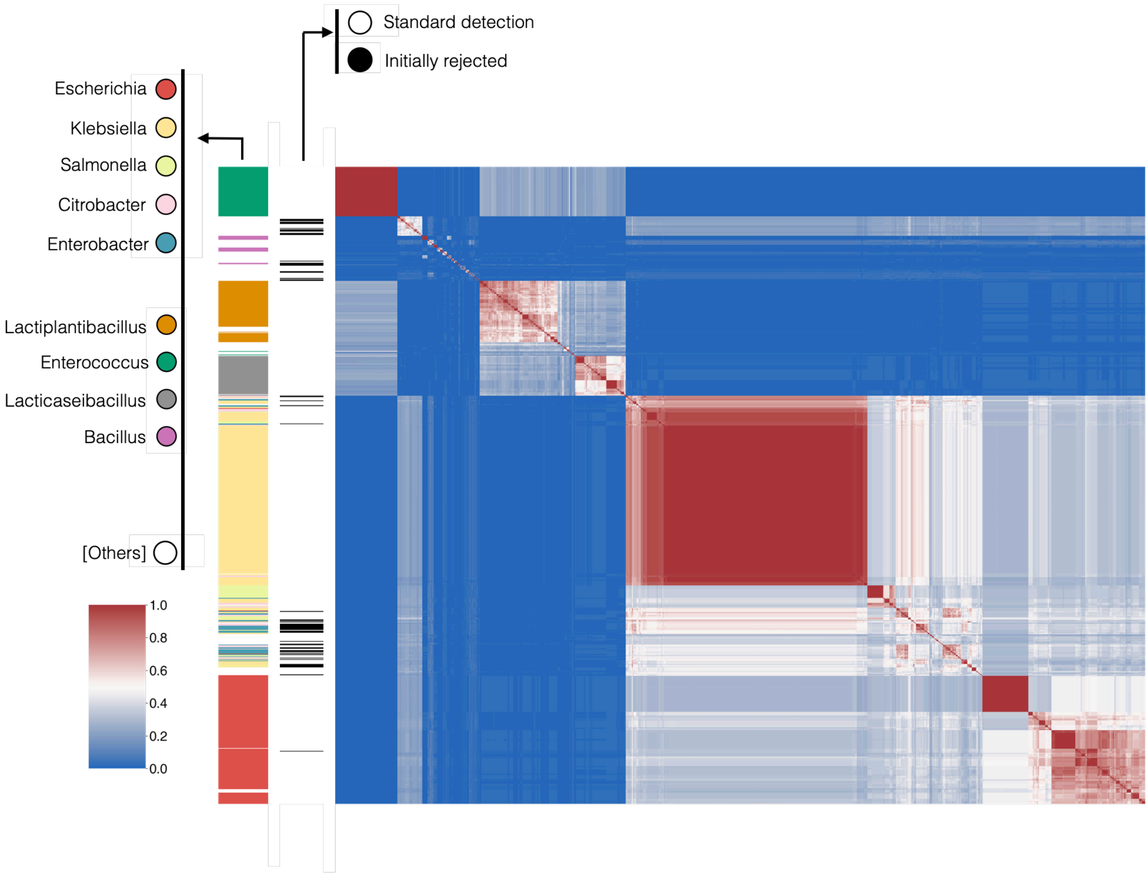

**Fig S8. Localization of initially rejected cfPICIs within the wGRR clusters of the family's genomes**

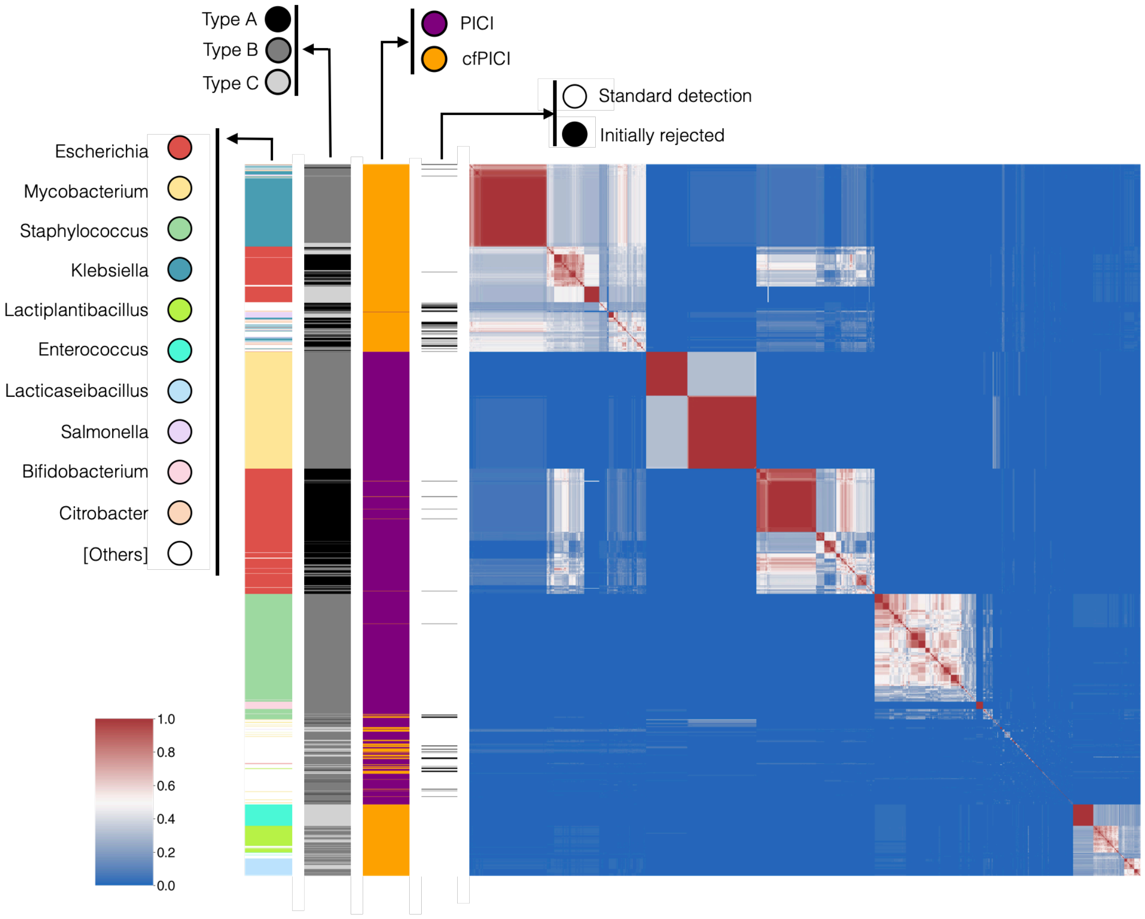

**Fig S9. wGRR comparison between PICI and cfPICI**

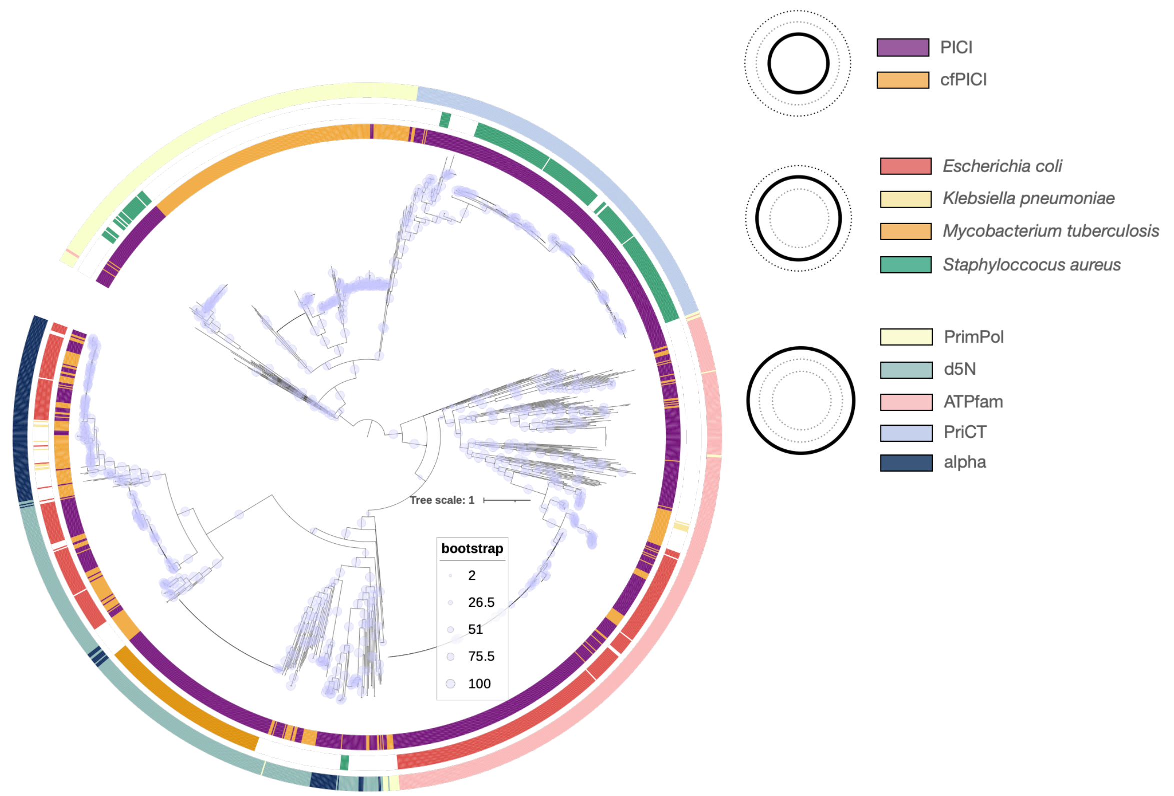

**Fig S10. Phylogenetic tree of the Primase-Replicase component in PICI and cfPICI**

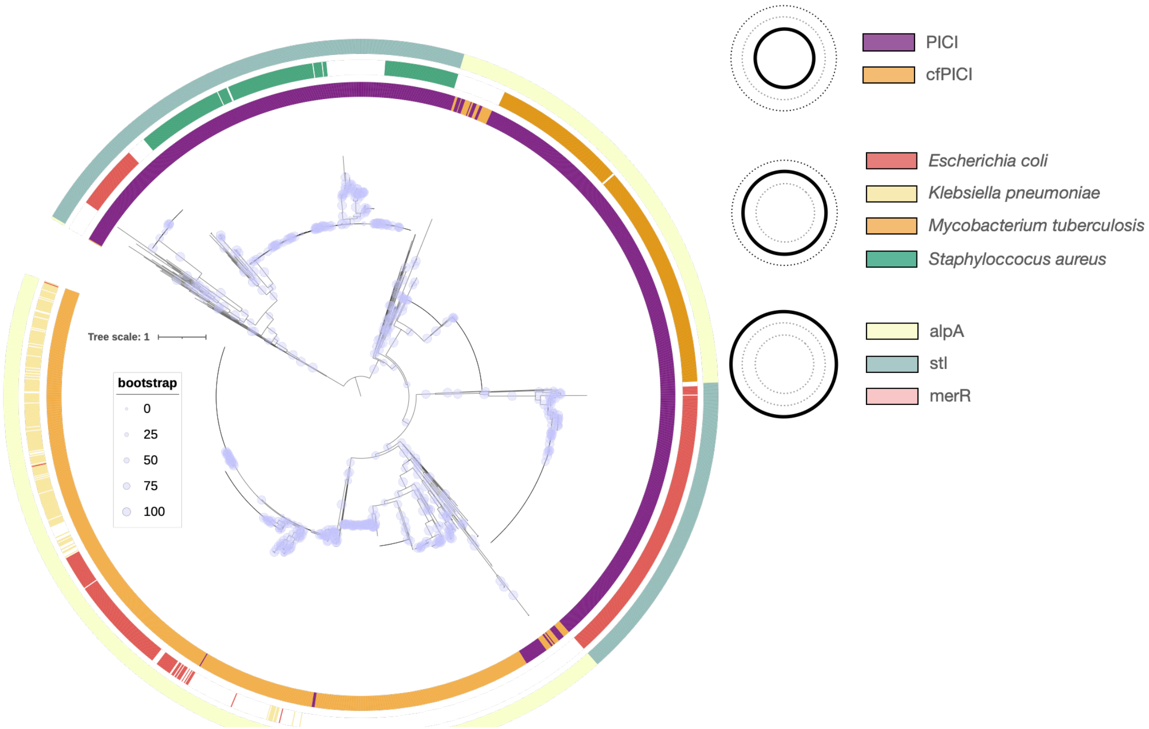

**Fig S11. Phylogenetic tree of the small terminase in PICI and cfPICI**

#### Table S1. SatelliteFinder classification of known phage-satellites

| Model Satellite | Classfied as | Type | Notes |
|---|---|---|---|
| PLE1 | PLE | Type A | |
| PLE2 | PLE | Type D | |
| PLE3 | PLE | Type F | |
| PLE4 | PLE | Type D | |
| PLE5 | PLE | Type B | |
| EcCISMS-3-5 | cfPICI | TypeB | |
| EcCIPNUSAE044409 | cfPICI | TypeC | |
| SeCIDerby | cfPICI | TypeB | |
| YaCI159 | cfPICI | TypeB | |
| SfCI301 | cfPICI | TypeA | |
| SfCI8401 | cfPICI | TypeA | |
| EcCIIAI39 | cfPICI | TypeA | |
| CfCICFNIH4 | cfPICI | TypeB | |
| EcCIEC11-7286 | cfPICI | TypeB | |
| EcCIED1a | cfPICI | TypeA | |
| EcCIWW223 | cfPICI | TypeB | |
| EpCIETW41 | cfPICI | TypeB | |
| PgCIFDAARGOS.186 | cfPICI | TypeA | |
| SdCICCFSAN010956 | cfPICI | TypeA | |
| SeCIFDA336426-1 | cfPICI | TypeA | |
| PmCIATCC29906 | cfPICI | TypeA | |
| EcCIEO709 | cfPICI | TypeB | |
| EcCIRM10386 | cfPICI | TypeA | |
| KvCIGJ3 | cfPICI | TypeA | |
| EcCICFSAN002236 | cfPICI | TypeA | |
| EcCID8 | cfPICI | TypeB | |
| EcCIRM10042 | cfPICI | TypeA | |
| CpCITV06 | cfPICI | TypeA | |
| EcCIHUST159 | cfPICI | TypeB | |
| SeCI08-1209 | cfPICI | TypeB | |
| EcCI392917 | cfPICI | TypeB | |
| SeCIKentucky | cfPICI | TypeB | |
| GaCISCGC | cfPICI | TypeB | |
| GaCIWKB1 | cfPICI | TypeA | |
| KpCIAR.0148 | cfPICI | TypeB | |
| EcCIEDL933 | cfPICI | TypeB | |
| EcCI144 | cfPICI | TypeA | |
| EcCIPA40 | cfPICI | TypeB | |
| SeCI7830 | cfPICI | TypeD | |
| EcCIST130 | cfPICI | TypeA | |
| KpCIKPN01 | cfPICI | TypeB | |
| EcCIPSUO103 | cfPICI | TypeB | |
| KoCICAV175 | cfPICI | TypeB | |
| XnCIATCC1906 | cfPICI | TypeA | |
| EcCI315650 | cfPICI | TypeA | |
| B..kocchi.BDGP4 | cfPICI | TypeB | |
| B.cereus.BAG6X1-2 | cfPICI | TypeB | |
| S.saprophyticus.CCUG38042 | cfPICI | TypeA | |
| C.botulinum.B.Eklund.17B | cfPICI | TypeF | Components are too apart, detected as two sets |
| C.botulinum.B.Eklund.17B | cfPICI | TypeG | Components are too apart, detected as two sets |
| S.pettenkoferi.589 | cfPICI | TypeB | |
| S.haemolitycus.S167 | cfPICI | TypeA | |
| S.equorum.DSM15097 | cfPICI | TypeA | |
| S.aureus.VET0180R | cfPICI | TypeA | |
| S.xylosus.HKUOPL8 | cfPICI | TypeA | |
| C.beijerinckii.WB53 | cfPICI | TypeB | |
| S.arlettae.IOV5 | cfPICI | TypeA | |
| S..aureus.C0673 | cfPICI | TypeA | |
| E.durans.4928STDY7071318 | cfPICI | TypeB | |
| S.warneri.DE0454 | cfPICI | TypeC | |
| L.rhammosus.GG | cfPICI | TypeB | |
| B.thuringiensis.BGSC.4W1.4W1 | cfPICI | TypeB | |

| | | |
|---|---|---|
| L.casei.Lc705 | cfPICI | TypeB |
| C.sporogenes.87-0535 | cfPICI | TypeA |
| S.hominis.SNUC.5746 | cfPICI | TypeA |
| L.casei.BL23 | cfPICI | TypeB |
| GN_EcCI11368.1 | PICI | TypeA |
| GN_EcCIIHE3034 | PICI | TypeA |
| GN_EcCIRM13514 | PICI | TypeC |
| GN_EcCIRM12579 | PICI | TypeA |
| GN_EcCIATCC_25922 | PICI | TypeA |
| GN_EcCI042 | PICI | TypeB |
| GN_PcCIPCC221 | PICI | TypeC |
| GN_EcCIDi14 | PICI | TypeA |
| GN_SbCISb277 | PICI | TypeB |
| GN_PmCIOH1905 | PICI | TypeC |
| GN_PhaCIATCC43949 | PICI | TypeC |
| GN_EcCICFT073 | PICI | TypeB |
| GN_EcCI11128 | PICI | TypeA |
| GP_SAPI1 | PICI | TypeB |
| GP_LlCIKF147 | PICI | TypeC |
| GP_LlCIA76-1 | PICI | TypeB |
| GP_SaPImw2 | PICI | TypeC |
| GP_SAPI2 | PICI | TypeB |
| GP_SpnCITaiwan-0.2 | PICI | TypeC |
| GP_SpnCITaiwan-0.03 | PICI | TypeC |
| GP_SpnCI-A45-1.9 | PICI | TypeC |
| GP_SaPIbov1 | PICI | TypeB |
| GP_LlCISK11 | PICI | TypeB |
| GP_SpnCITCH8341-0.25 | PICI | TypeC |
| GP_SpnCIINV104-1.06 | PICI | TypeC |
| GP_EfCIV583 | PICI | TypeC |
| GP_MG1363-1 | PICI | TypeC |
| GP_LlCI-CV56-1 | PICI | TypeB |
